# Investigating the Conformational Ensembles of Intrinsically-Disordered Proteins with a Simple Physics-Based Model

**DOI:** 10.1101/2020.02.11.943969

**Authors:** Yani Zhao, Robinson Cortes-Huerto, Kurt Kremer, Joseph F. Rudzinski

## Abstract

Intrinsically disordered proteins (IDPs) play an important role in an array of biological processes but present a number of fundamental challenges for computational modeling. Recently, simple polymer models have re-gained popularity for interpreting the experimental characterization of IDPs. Homopolymer theory provides a strong foundation for understanding generic features of phenomena ranging from single-chain conformational dynamics to the properties of entangled polymer melts, but is difficult to extend to the copolymer context. This challenge is magnified for proteins due to the variety of competing interactions and large deviations in side-chain properties. In this work, we apply a simple physics-based coarse-grained model for describing largely disordered conformational ensembles of peptides, based on the premise that sampling sterically-forbidden conformations can compromise the faithful description of both static and dynamical properties. The Hamiltonian of the employed model can be easily adjusted to investigate the impact of distinct interactions and sequence specificity on the randomness of the resulting conformational ensemble. In particular, starting with a bead-spring-like model and then adding more detailed interactions one by one, we construct a hierarchical set of models and perform a detailed comparison of their properties. Our analysis clarifies the role of generic attractions, electrostatics and side-chain sterics, while providing a foundation for developing efficient models for IDPs that retain an accurate description of the hierarchy of conformational dynamics, which is nontrivially influenced by interactions with surrounding proteins and solvent molecules.

## I. Introduction

Despite lacking stable tertiary structure under physiological conditions, intrinsically disordered proteins (IDPs) are involved in a large number of important biological functions, including intracellular signaling and regulation, and are also associated with a broad range of diseases, including cancer, neurodegenerative diseases, amylidoses, diabetes and cardiovascular disease^1,2^. The experimental characterization of IDPs is complicated by the heterogeneous nature of their disordered conformational ensembles (i.e., conformational distributions), which challenges traditional techniques developed for folded proteins. For example, X-ray crystallography and cryo-EM, which recover high resolution images of biomolecules in the crystalline or frozen state, are fundamentally inappropriate for characterizing the distribution of relevant IDP conformations^3^. On the other hand, techniques including Nuclear Magnetic Resonance (NMR), Small Angle X-ray Scattering (SAXS), single-molecule Förster resonance energy transfer (FRET), dynamic light scattering (DLS) and two-focus fluorescence correlation spectroscopy (2f-FCS) are capable of identifying the conformational transitions sampled by IDPs^4–7^, since they perform measurements of the protein as it fluctuates within its “natural” environment. However, these measurements provide limited resolution in terms of the specification of a unique corresponding microscopic distribution of conformations. In other words, there may exist multiple distinct conformational ensembles which reproduce the experimental measurements, requiring molecular models to infer the correct underlying distribution. As a result, molecular simulations have become increasingly important tools for obtaining microscopic insight that supports experimental observations, e.g., for the characterization of IDP conformational ensembles^4^.

All-atom (AA) models have gained significant popularity for providing detailed descriptions of complex biomolecular processes and, in conjunction with reweighting techniques, can also be used to assist in the interpretation of experimental measurements. The application of AA simulations to study IDPs has brought to light transferability problems of standard models, which were constructed to stabilize three-dimensional structures of folded proteins. Not only do these force fields predict overly compact structures^8^, distinct AA models can generate widely varying and qualitatively different secondary structure content for a given protein sequence^9^. Recent efforts have been made to adjust these models to more accurately describe the properties of IDPs^8,10,11^. Despite these improvements, AA simulations remain prohibitively expensive for investigating the environment-dependent conformational dynamics of IDPs, due to the expansive conformational landscape traversed by these systems. Moreover, the large range of time scales (from ps to hours), thermodynamic or chemical conditions (e.g., denaturation concentrations), as well as system variations (e.g., sequence mutations) commonly explored in experimental studies represents an overwhelming gap in computational accessibility for AA models that is unlikely to be overcome in the near future through improvements in software or hardware.

The computational expense of these detailed models has motivated the use of much simpler polymer models^12,13^, e.g., ensemble construction methods^14^ or analytically-solvable polymer models^4^, to provide microscopic interpretations for the experimental characterizations of processes involving IDPs. The disordered nature of IDPs results in conformational heterogeneity and broad intramolecular distance distributions, reminiscent of generic models from the study of polymer physics^15^. However, these models are limited in resolution, and often lack the ability to provide significant microscopic insight beyond what can already be inferred from experiments. Moreover, the simplicity of the model approximations have been shown to generate inconsistencies in the interpretation of experimental measurements.^16–19^ Native-biased models, e.g., Gō-type models,^20^ which use experimentally-determined protein structures to construct a potential energy function with the protein’s native state at the global minimum, have contributed immensely to our basic understanding of the driving forces for protein folding.^21–23^ When combined with additional non-native interactions, these models provide a straightforward route to elucidate the essential features for reproducing a given experimental observation.^24–26^ Although these models have been useful for investigating the environment-dependent folding processes of IDPs^27,28^ (i.e., coupled folding and binding processes), their reliance on a well-defined native structure limits their ability to describe unfolded or disordered conformations. This limitation can even propagate into the characterization of the folding process of globular proteins, resulting in a qualitatively incorrect representation of folding pathways^29^. Recent work from Shell and coworkers aims to partially alleviate this limitation by combining transferable bonded interactions with traditional native-like “nonbonded” interactions^30^. There have also been significant advancements in the development of physics-based CG models to describe the temperature-dependent collapse and liquid-liquid phase separation of IDPs^31,32^.

Recently, Rudzinski and Bereau proposed a simple physics-based model^33^ for describing largely disordered conformational ensembles of peptides. The foundational premise of the model is that the sampling of sterically-forbidden conformations, due to missing degrees of freedom, can seriously complicate the faithful description of both static and dynamic properties in coarse-grained (CG) models of proteins. This complication is perhaps most severe for disordered ensembles, where conformational entropy plays an important role in shaping the free-energy landscape. For this reason, the steric interactions and local stiffness of the protein are described at a united-atom resolution (i.e., explicit representation of all heavy atoms). These interactions account only for excluded volume and chain stiffness, without explicit attractions between atoms which reside at significant separation along the peptide chain. In addition to these detailed interactions, coarse-grained attractive interactions are added to represent the characteristic driving forces for peptide secondary and tertiary structure formation. For example, in the introductory studies^33,34^, the authors employed a generic attractive interaction between *C*_*β*_ carbons in order to model the effective attractions between side chains due to the hydrophobic effect. Additionally, attractive interactions between *C*_*α*_ atoms separated by three peptide bonds along the protein backbone were employed to model helix-forming hydrogen-bonding interactions. These two interactions represent the minimum set of interactions necessary for qualitative reproduction of the conformational ensemble of short peptides, i.e., to sample helical, coil, and swollen (i.e., hairpin-like) structures. The model was shown to accurately characterize both structural and kinetic properties of helix-coil transitions in small peptides, demonstrating its potential for efficiently describing disordered ensembles, while retaining relevant microscopic details^33,34^. Furthermore, the Hamiltonian of the model can be easily adjusted to investigate the driving forces for particular processes.

In this work, we apply variants of this simple physics-based model to investigate the role of distinct interactions in shaping disordered protein ensembles. As a model system, we consider the activation domain, ACTR, of the SRC-3 protein, a “fully disordered” protein with only transient helical propensity^35–37^. One way in which IDPs perform their function is by adapting to their environment through so-called coupled folding and binding processes^38^. For example, ACTR can form a structured complex with the nuclear-coactivator binding domain (NCBD) of the transcriptional coactivator CREB-binding protein (CBP), which plays an important role in the regulation of eukaryotic transcription^39^. CBP demonstrates the functional advantages of IDPs in the regulation of genes^39^, participating in interactions with more than 400 transcription factors in the cell^40^. In the absence of a binding partner, NCBD is a molten globule with three substantial helical regions^39^, but undergoes coupled folding and binding processes with a variety of distinct ligands^41^. Within the NCBD/ACTR complex, the three helices of NCBD form a bundle with a hydrophobic groove in which ACTR is docked, and the assembly of the two proteins promotes three helices in ACTR^37^ (see Figure 1).

**FIG. 1:**
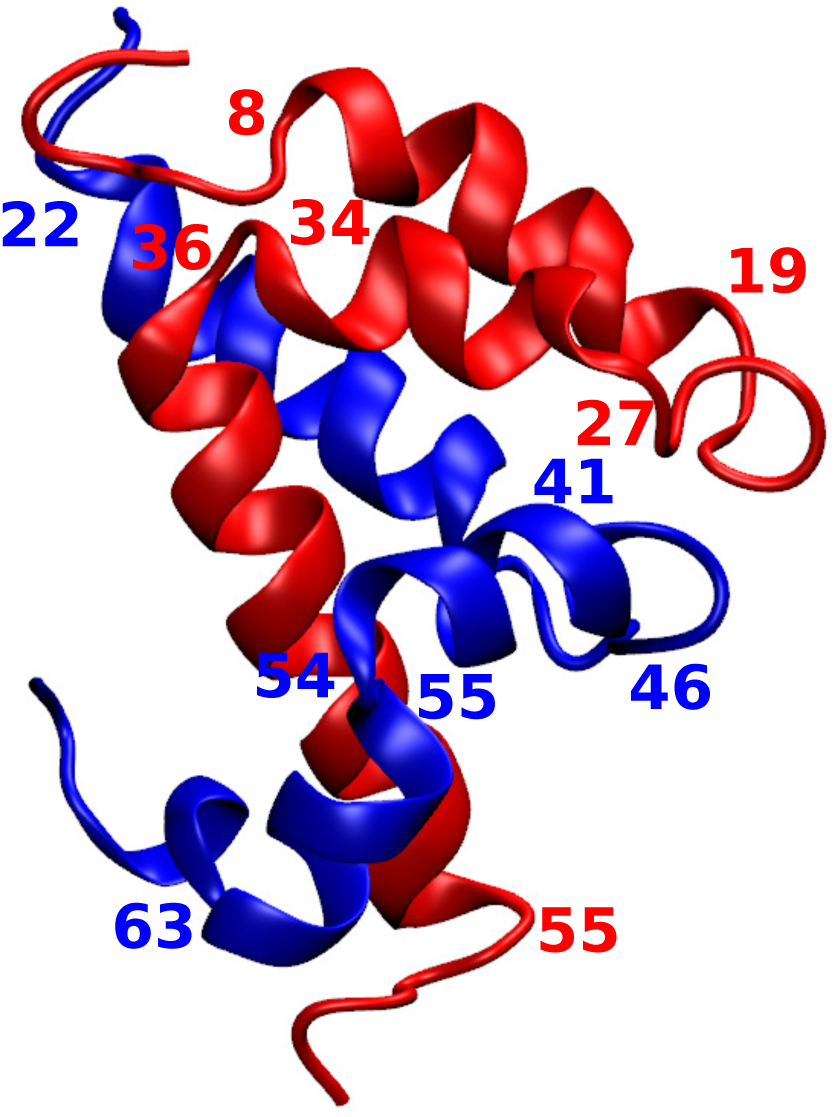
Visualization of the NCBD/ACTR folded complex (PDB-id: 1KBH). The number labels correspond to residue numbers at the beginning and ends of helices formed by NCBD (red) and ACTR (blue).

Great efforts have been made to understand how IDPs recognize their binding partners^42,43^. For example, electrostatic attractions have been shown to play an important role in driving the formation of encounter complexes between the binding pair^38,44^. Additionally, the change in solvent accessible surface area of IDP residues upon binding suggests that IDPs can utilize different residues along the amino acid sequence for interactions with different binding partners^2^. The conformational diversity of IDPs leading to the folded state makes it challenging to precisely characterize their binding mechanisms both experimentally and computationally^45,46^. Previous work has identified two limiting mechanisms—”conformational selection” and “induced fit”— of coupled folding and binding, although in practice a combination of these is typically observed^35,47,48^. The conformational selection mechanism is characterized by an IDP which samples the relevant folded structure (or some fraction of this structure) within the unbound ensemble. In the induced fit mechanism, the folded state only arises within the conformational ensemble of the IDP through interactions with its binding partner. Thus, a key step to describing the binding mechanism for a particular IDP/partner pair, especially in cases where conformational selection is prominent, is to characterize the unbound ensembles of the molecules. Previous computational work employing Gō-type models has found that the NCBD/ACTR folding process demonstrates dominant characteristics of the induced fit mechanism^47^. However, a mechanistic shift toward conformational selection is also possible when NCBD attains a distinct folded structure after a proline isomerization^49^. Computational investigations of coupled folding and binding typically employ models that are not explicitly constructed to accurately model the unbound ensembles of the individual binding partners. While the unbound ensemble of NCBD has been analyzed using both AA and CG simulations^35,36,50^, the unbound conformational behavior of ACTR has not, to our knowledge, been investigated in detail. Instead, ACTR is usually taken to be a fully-disordered ensemble, as characterized by simple polymer models^15,51^.

The present investigation employs an intermediate resolution physics-based model to characterize the ensemble of ACTR in the absence of a binding partner. This model enables systematic analysis of the impact of distinct interactions and sequence specificity on the randomness of the resulting conformational ensemble. In particular, starting with a bead-spring-like model and then adding more detailed interactions one by one, we construct a hierarchical set of models and perform a detailed comparison of their properties. Our analysis shows that: (i) the incorporation of generic attractions between amino acid side chains significantly expands the diversity of the conformational ensemble, without severely perturbing the distribution of the radius of gyration, (ii) electrostatic interactions can increase the ruggedness of the conformational landscape, reducing the overall conformational heterogeneity, but can simultaneously introduce additional routes for stabilizing particular secondary structures, and (iii) side chain sterics play a crucial role in determining the overall shape of the free-energy landscape through stabilization of particular structural motifs.

The rest of the paper is organized as the follows. In section II, the hierarchical set of models, associated simulation protocol, and relevant analysis tools are described in detail. Section III presents a detailed characterization of the hierarchy of CG models describing the unbound conformational ensemble of ACTR. Two additional polypeptides are also considered to investigate the effect of side chain excluded volume on the conformational ensembles of IDPs. Then, the transferability to the unbound ensemble of NCBD using the more detailed models is assessed. Finally, Section IV provides a brief discussion and conclusions from the investigation.

## II. Methods

### A. Protein sequences

This work considers the activation domain, ACTR, of the SRC-3 protein and the nuclear-coactivator binding domain (NCBD) of the transcriptional coactivator CREB-binding protein (CBP). The amino acid sequences of ACTR and NCBD are given by:

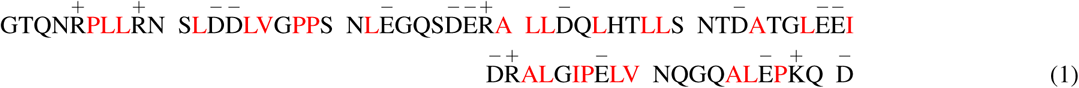

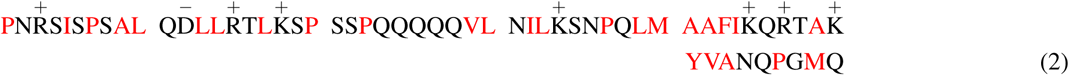

where Equations (1) and (2) are the sequences for ACTR and NCBD with 71 and 59 residues, respectively. The spacing in the equations separate the sequences into groups of 10 residues. Hydrophobic residues are labeled with red font, while the positively and negatively charged residues are denoted by “+” and “-”, respectively. Upon interaction, NCBD and ACTR form a stable folded complex (PDB-id: 1KBH, see Figure 1).

### B. A simple physics-based model for describing disordered ensembles

ACTR and NCBD were modeled using a physics-based approach that represents the protein in near-atomic detail while treating the solvent implicitly through effective interactions between protein atoms^33,34^. The total potential energy function of the model can be written as a sum of three terms:

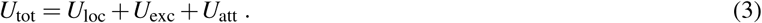

*U*_loc_ represents local interactions contributing to chain connectivity and stiffness and employs the standard functional forms and parameters for bond, angle, dihedral, and 1-4 interactions given by the Amber99SB-ILDN force field^52,53^ (see Figure S1 in the Supporting Information). For reasons that will become clear below, we write *U*_loc_ as a sum of two contributions:

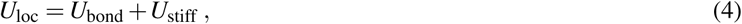

where *U*_bond_ represents the bond interactions between pairs of covalently bonded atoms and *U*_stiff_ represents the remaining local interactions listed above. *U*_exc_ represents excluded volume interactions at a united-atom resolution (i.e., an explicit representation of all heavy atoms, without hydrogens). The excluded volume interaction for each heavy atom pair was determined by transforming the Lennard-Jones interactions between the pair (again given by the Amber99SB-ILDN force field) to a Weeks-Chandler-Andersen (WCA) potential (i.e., a purely repulsive potential). *U*_att_ represents the attractive interactions employed between *C*_*α*_, *C*_*β*_, and representative side-chain atoms and can contain several distinct contributions. In this work, we consider a hierarchy of eight different models which systematically build upon each other (see Table I). The first model employs only bond and excluded volume interactions, i.e., *U*^(1)^ = *U*_bond_ + *U*_exc_, similar to the self-avoiding random walk model from polymer theory^12^. The second model adds stiffness to the chain by incorporating the other local interactions: *U*^(2)^ = *U*_loc_ + *U*_exc_.

**TABLE I:** Overview of interactions for model hierarchy.

| model id | $U_{\text{bond}}$ | $U_{\text{stiff}}$ | $U_{\text{exc}}$ | $U_{\text{hp}}^{(x)}$ | hp type | $U_{\text{hb}}$ | $U_{\text{DH}}$ |
| --- | --- | --- | --- | --- | --- | --- | --- |
| 1 | YES | NO | YES | NO | N/A | NO | NO |
| 2 | YES | YES | YES | NO | N/A | NO | NO |
| 3a | YES | YES | YES | YES | HOMO | NO | NO |
| 3b | YES | YES | YES | YES | HETERO | NO | NO |
| 4 | YES | YES | YES | YES | HETERO | NO | YES |
| 5a | YES | YES | YES | YES | HOMO | YES | NO |
| 5b | YES | YES | YES | YES | HETERO | YES | NO |
| 6 | YES | YES | YES | YES | HETERO | YES | YES |

The remaining models employ the full *U*_tot_ potential with varying representations of *U*_att_, i.e., 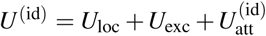, for id ∈ {3a,3b,4,5a,5b,6}. Model 3 employs attractive interactions between *C*_*β*_ atoms, *U*_hp_, to model the hydrophobic attraction between side chains: 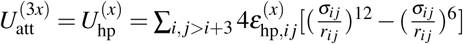 with *σ*_*i j*_ =0.5 nm. We consider two variants of model 3: (i) *x* = *a*, where the same parameter is employed for all amino acids (denoted homo), 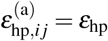 and (ii) *x* = *b*, where the parameter depends on the identity of the pair of residues (denoted hetero), 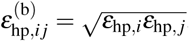, where *ε*_hp,*i*_ is determined according to the Miyazawa-Jernigan interaction matrix^54^ (see Figure S2 and Table S1). More specifically, to set the absolute scale of these interactions, we followed the work of Bereau and Deserno^55^. Briefly, the 20×20 Miyazawa-Jernigan interaction matrix is reduced to 20 residue-specific energy values, which approximately generate the full matrix through geometric averages between pairs of residue types. These energy values are then normalized to be between 0 (most hydrophilic) and 1 (most hydrophobic). Finally, a single overall interaction scale, *ε*_hp_, is chosen to determine all values of *ε*_hp,*i*_ simultaneously. Model 4 builds upon model 3 by incorporating electrostatic interactions, *U*_DH_, between charged residues: 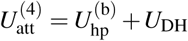. These interactions are described at a coarse-grained level of resolution (see Figure S3), using the Debye-Hückel formalism^56^, where the full point charge is placed on the last side chain carbon (i.e., furthest from the backbone) for each charged residue, i.e., arginine (R), lysine (K), aspartic acid (D), and glutamic acid (E). In particular, the electrostatic energy is given by

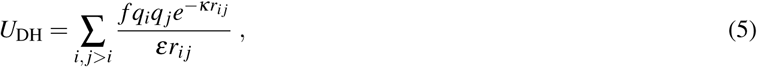

where 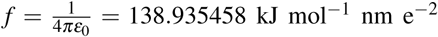 and *ε* = 80 at room temperature for monovalent salt. *κ*^−1^ is the Debye screening length, *q*_*i*_ and *q* _*j*_ are the point charges of the *i*^*th*^ and *j*^*th*^ charged sites, and *r*_*i j*_ is the distance between these sites. *κ*^−1^ = 0.313 *I*^−1*/*2^ nm mol^1*/*2^ L^−1*/*2^, where 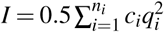 is the ionic concentration of the solution, *n*_*i*_ is the number of unique ionic species and *c*_*i*_ is the molar concentration of the ion type *i* with charge *q*_*i*_^57^. Employing physiological concentrations *c*_*i*_ = 0.1 mol/L for all ions, we obtain *κ*^−1^ = 1 nm.

Model 5 builds upon model 3 by incorporating “local” hydrogen-bonding interactions, *U*_hb_, between *C*_*α*_ atoms that are separated by three residues along the peptide backbone: 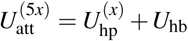. This interaction ensures that the proteins are capable of forming *α*-helical conformations. The incorporation of 1-4 hydrogen bonds independently from hydrogen bonds occurring between residues farther apart along the peptide chain allows the independent investigation of the driving forces for helical versus *β* -sheet conformations. The latter are not considered in the present study since ACTR does not have a substantial propensity toward *β* -sheet formation. In a way, the local hydrogen bonds represent a “native-like” interaction for peptides that fold into a single helix. For this reason, the model was originally designated as a “hybrid Gō” model, indicating the combination of atomically-detailed physics-based interactions with simplistic (possibly natively-biased) attractive interactions at a coarser level of resolution. Note that in previous work the hydrogen-bonding interaction was denoted nc for “native contact”. Following previous work employing native-biased CG models, we employ a hydrogen-bonding interaction with a Lennard-Jones form along with a desolvation barrier using the following functional form^24^: 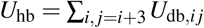, where

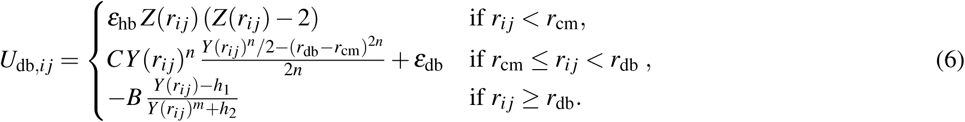

In Equation (6), *r*_cm_ = 0.5 nm is the position of the first potential minimum with a corresponding depth of *ε*_hb_, and *r*_db_ = 0.65 nm is the position of the desolvation barrier maximum with a corresponding height of *ε*_db_ = 0.4*ε*_hb_. 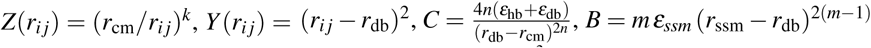 with *ε*_ssm_ = *ε*_db_/100 and 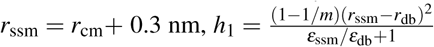 and 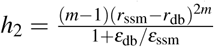. The parameters *k* = 6, *m* = 3 and *n* = 2 control the shape of *U*_hb_ (See^33^ for a plot of the potential). Again, two variants of 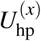 are considered, with homo- and hetero-type interactions for *x* = a and *x* = b, respectively, as described above. Finally, model 6 also incorporates electrostatic interactions: 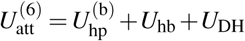. The hierarchy of models employed in this work is summarized in Table 1.

Previous work using model 4 performed an extensive search in parameter space to characterize the behavior of the model in the context of helix-coil transitions of short peptides^33,34^. Here, we tune the parameters of the model in an attempt to accurately describe the conformational ensemble of ACTR. There are no adjustable parameters for the local, excluded volume, and electrostatic interactions. Moreover, as described above, several of the parameters for the hydrogen-bonding interactions have been fixed based on previous work^33,34^. Thus, the models are left with just two free parameters: *ε*_hp_ and *ε*_hb_. *ε*_hp_ was initially determined by simulating model 3a with various parameter values and then comparing the generated *R*_*g*_ distribution with that determined from experimental measurements^6^. For model 3b (hetero hp type), the residue-specific hydrophobic attractions were applied such that the average hydrophobic interaction energy (i.e., the average value of *ε*_hp,*i*_ along the chain) was identical to that of model 3a (homo hp type). 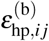 for ACTR is presented in the Supporting Information (Figure S3). After fixing *ε*_hp_, *ε*_hb_ was determined by simulating model 4 with various parameter values and then comparing the generated average fraction of helical segments per residue, *h*(*i*), to experiments^6^. With the exception of the difference in *ε*_hp_ used for the homo and hetero variants, described above, identical *ε*_hp_ and *ε*_hb_ parameters were employed for the entire hierarchy of models, wherever applicable. We hypothesize that the very accurate representation of sterics in the model will result in energetic parameters that are quite sequence transferable, for sequences that exhibit largely disordered ensembles. A challenging test of transferability is assessed toward the end of this work by considering the molten globule NCBD.

Note that when comparing models with fundamentally different interactions, there is no unique procedure for calibrating the energy scales of the models. When the interaction sets are not entirely different (as is the case for the hierarchy of models considered here), one option is to evaluate the models on the same absolute temperature scale, as dictated by the simulation protocol. This would lead to different ensemble properties at the relevant temperature, due to changes in the incorporated interactions. Alternatively, one can work with a reduced temperature scale, defined with respect to a reference temperature, *T*^∗^, at which a particular ensemble property is reproduced. We follow this latter approach in the present work, and define *T*^∗^ as the temperature at which the average experimental radius of gyration is reproduced. In terms of the absolute temperatures employed in the simulation protocol, *T*^∗^ corresponds to 300 K for ACTR for models 3a, 3b, 5a and 5b, and 270 K for models 4 and 6. For NCBD, *T*^∗^ corresponds to 330 K for the two considered models, 5b and 6.

### C. Simulations

All simulations of the hierarchical set of physics-based models were performed with the GROMACS 4.5.5 simulation suite^58^ in the constant *NVT* ensemble while employing the stochastic dynamics algorithm with a friction coefficient *γ* = (2.0 *τ*)^−1^ and a time step of 1 × 10^−3^ *τ*. The CG unit of time, *τ*, can be determined from the fundamental units of length, mass, and energy of the simulation model. Employing any one of the Lennard-Jones radii and energies from the Amber99SB-ILDN force field yields a time unit on the order of 1 ps. We report the connection to physical units since the models are simulated using these units within the GROMACS suite. For simplicity, we define *τ* = 1 ps, and report the simulation protocol in units of *τ*. Note that this relationship to physical units does not provide any meaningful description of the absolute time scale of characteristic dynamical processes generated by the model, due to a lost connection to the true dynamics^59^. The present study focuses on ensemble-averaged properties of the generated ensembles, and does not attempt to calibrate or interpret the generated dynamics, although previous studies with this model have demonstrated the faithful reproduction of kinetic processes for secondary-structure formation^33,34^. For each peptide, a single chain was placed in a cubic box with a volume of (20 nm)^3^ and simulated *without* periodic boundary conditions. Thus, no explicit cutoffs were used for the interaction functions described in the previous section. Replica exchange simulations^60^ were performed to enhance the sampling of the system. In total, 16 temperatures ranging from 225 to 450 K were scanned with an average acceptance ratio of 0.4. Note that these represent absolute simulation temperatures, which were transformed to reduced temperatures for comparison of different models (as described above). The exchange of replicas was attempted every 500 or 1,000 *τ*, and each simulation was run for at least 500,000 *τ*. The convergence of the simulations were assessed by randomly dividing each trajectory into two groups and then checking for consistency of various observables, including the average radius of gyration and the average fraction of helical segments, as well as autocorrelation functions of the radius of gyration and of the end-to-end distance. Representative examples of the convergence tests are presented in Figures S4 and S5.

For comparison with more generic polymer ensembles, we considered a bead-spring (BS) model (often referred to as the Kremer-Grest model)^61^, which represents each monomer (i.e., residue) with a single coarse-grained site. Connections between monomers are represented with the finite extensible nonlinear elastic (FENE) potential. We considered two variations of the BS model, which differed in the treatment of nonbonded interactions. The first (denoted “BS”) employed a purely repulsive WCA potential to represent interactions between monomers, while the second (denoted “BS-LJ”) employed a standard Lennard-Jones (LJ) potential with a cutoff *r*_*c*_ = 2.5*σ*. The properties of the BS model are determined in reduced units in terms of the LJ interaction radius, *σ*, the well depth, *ε*, and the mass, *m*, of a monomer. The corresponding time unit is 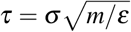. The BS models were simulated at a temperature of *T*^∗^ = 2.0 *ε/k*_*B*_. Simulations of the BS model were performed with the ESPResSo++ package^62^. Each simulation employed a time step of 0.005 *τ* and was run for 3.2 × 10^9^ *τ*, while using the Langevin thermostat with a damping coefficient of 1.0 *τ*^−1^.

### D. Analysis

#### Polymeric behavior

Because IDPs possess some similar properties to more generic polymer systems, such as long-range fluctuations and structural heterogeneity, traditional polymer physics analysis can be useful for providing an overarching description of the conformational ensembles of IDPs^15^. The single-chain backbone structure factor, which characterizes the overall shape of a molecule, is given by^63,64^:

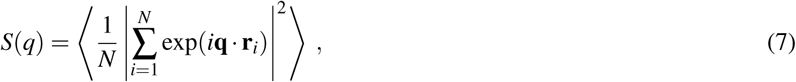

where *N* is the number of residues (*N* = 71 for ACTR and *N* = 59 for NCBD) and **q** is the wave vector. **r**_*i*_ corresponds to the position of the C_*α*_ atom of the *i*^*th*^ residue for the physics-based models and the position of the *i*^*th*^ bead for the BS models. *S*(*q*) is widely used to characterize polymer systems^64^. We also calculated the shape parameters 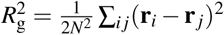 (radius of gyration), 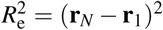 (end-to-end distance), and 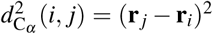 (inter-residue distance between the C_*α*_ atoms). We will use the notation 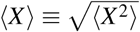, where 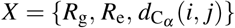. The average (real space) distance between two residues separated by *m* residues along the chain is calculated as 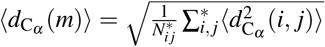, where 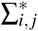 is a sum over all *i j* pairs with | *j* − *i* |= *m* and 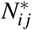 is the number of such pairs. Note that 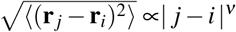, where *v* is the Flory scaling exponent. Thus, it is useful to consider the normalized quantity 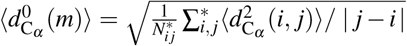, such that 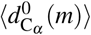 is constant for a random walk and proportional to *m*^0.1^ for a self-avoiding random walk^13,64^.

For a slightly more detailed view of the ensemble, we also calculated contact probability maps, which are obtained by determining the probability that a pair of C_*α*_ atoms are within a given cutoff distance, *r*_c_, from one another. In this case, we have chosen *r*_c_ = 1.0 nm. Additionally, we calculated the gyration tensor 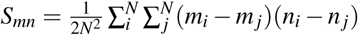, where *m, n* ∈ {*x, y, z*}. Note that only the C_*α*_ atoms were taken into account in the calculation of the gyration tensor, for consistency with the BS models. The eigenvalues of *S*_*mn*_ are calculated and ordered as *λ*_1_ ≤ *λ*_2_ ≤ *λ*_3_. The asphericity of the chain can be characterized in terms of these eigenvalues: 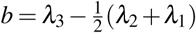. The asphericity values reported throughout the text are normalized by 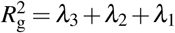: 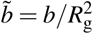. For a self-avoiding random walk, the ratio of eigenvalues is *λ*_3_ : *λ*_2_ : *λ*_1_ ≅ 12 : 3 : 1, i.e., *λ*_3_*/λ*_1_ = 12 and *λ*_3_*/λ*_2_ = 465.

#### Helical propensity

The helical propensity of the peptide is characterized by the average fraction of helical segments, *h*(*i*), for each residue *i. h*(*i*) is calculated within the context of the Lifson-Roig formulation^66^, which represents the state of each residue as being in either a helical, h, or coil, c, state^67^. More specifically, *h*(*i*) is defined as the average propensity of sequential triplets of h states along the peptide chain. Following previous work^68^, we define the helical region of the Ramachandran (*ϕ, ψ*) map as *ϕ* ∈ [−160°, −20°] and *ψ* ∈ [−120°, 50°], although the precise definition has little impact on *h*(*i*).

#### Dimensionality reduction and clustering

The conformational landscape of disordered proteins is difficult to characterize within a low-dimensional representation. Linear dimensionality reduction methods typically fail to provide meaningful representations, due to the high level of structural heterogeneity and subtle distinctions between different subensembles. Nonlinear manifold learning methods overcome the limited ability of linear methods to capture nonlinear relationships in the data and can determine the low-dimensional embedding based on a wide variety of criteria. These methods have been more successful in finding low-dimensional embeddings which provide a clear picture of distinct structures in disordered landscapes^69,70^. Here we employed UMAP (Uniform Manifold Approximation and Projection), a type of multidimensional-scaling algorithm that attempts to find a balance between resolving global and local properties of the conformational landscape^71^. More specifically, given a set of *N* input features (e.g., intramolecular coordinates), the conformation of the peptide is defined within an *N*-dimensional space. UMAP obtains the optimal (nonlinear) projection into an *n*-dimensional space (*n* < *N*) using a cost function which simultaneously incorporates pairwise distances between conformations at the largest (global) and smallest (local) scales. In other words, the projection attempts to preserve these two sets of high-dimensional pairwise distances in the low-dimensional space, which results in the preservation of certain features of the conformational landscape. As input features, we employed pairwise distances between *C*_*α*_ atoms and angles between triplets of *C*_*α*_ atoms. To reduce the dimension of the input, we applied the following coarse-graining procedure. We divided the peptide into 4-residue segments and computed the minimum distance between atoms belonging to pairs of segments. Pairs of segments separated by less than 3 other segments were excluded. Thus, a total of 28 pairwise distances were included in the input features. We then applied the same segment representation to calculate the average angles between triplets of segments, again excluding any combinations where any pair of segments is separated by less than 3 other segments. This yields a total of 84 angles.

We performed UMAP with an embedding dimension of 2, using the standard Euclidean distance as the metric for evaluating similarity of structures (according to their input features). UMAP requires the choice of two other hyperparameters: the number of neighbors and the minimum distance. Over the range of hyperparameters considered, the resulting embedding space appeared to be relatively robust, but displayed a noticeable change in the “clustering” of data points as a function of either of the hyperparameters. We chose parameter values which resulted in “reasonable” clustering, i.e., a balance between a single cluster and a very diffuse landscape of points: 819 neighbors and 0.01 minimum distance. Since the conclusions made from this analysis is largely qualitative, we do not believe that the hyperparameter choice plays a significant role in our analysis. The UMAP projection was determined using the conformational ensemble generated by model 4. Subsequently, this projection was applied to the ensembles from each of the other models for consistent comparisons. This projection involves a “small” statistical component which has been shown to be normally distributed. Thus, we performed the projection 10 times for each configuration while randomly shuffling the input features. The average of the resulting UMAP coordinates were taken as the “true” projection and used to generate the free-energy landscapes presented below.

While nonlinear dimensionality reduction is necessary for providing a clear description of the overall conformational ensemble, linear methods are very effective if one is only interested in distinguishing between different helical states. Thus, we also applied principal component analysis on the conformation space characterized by the *ϕ/ψ* dihedral angles of each residue along the peptide backbone^72^. We then performed a k-means clustering^73^ along the largest three principal components in order to partition the conformation space into 50 states. We subsequently grouped these 50 *microstates* into 8 coarse-grained states by applying the PCCA+ dynamical coarse-graining method^74^.

## III. Results and Discussion

In this work, we characterize the role of distinct interactions in determining the disordered ensembles of IDPs. The focus of the study is the “fully disordered” peptide ACTR, which displays only transient helical structures. ACTR has 71 residues consisting of 26 hydrophobic residues, 18 charged residues, and a net charge of −8 (Equation 1). The average radius of gyration, 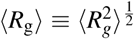, of ACTR, determined from small angle X-ray scattering experiments, is 26.5 Å at 5°C and 23.9 Å at 45°C. Note that the average size of ACTR decreases when the temperature is increased from 5°C to 45°C. It has been argued that many disordered proteins undergo such a collapse with increasing temperature due to the unfavorable solvation free energy of individual residues^75^. The temperature-dependent collapse of IDPs can be captured by atomistic simulations with explicit solvent, while temperature-dependent force field parameters are required for implicit solvent CG models^76^. For this reason, the present study focuses on the ensemble of conformations sampled at a single temperature. In particular, we focus on the higher temperature ensemble of ACTR, and investigate models which approximately reproduce 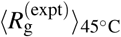. We employ an intermediate-resolution physics-based coarse-grained model, which represents the excluded volume of the peptide with united-atom resolution, while treating the attractive interactions which stabilize secondary and tertiary structure in a much coarser manner. The model also represents the solvent implicitly through these attractive interactions. We consider eight distinct models with different interaction sets, as summarized in Table 1 and described in detail in the Methods section. The models are separated into three groups: (i) models 1 and 2, without explicit attractive interactions, (ii) models 3a, 3b and 4, without hydrogen-bonding-like interactions, and (iii) models 5a, 5b and 6, with hydrogen-bonding-like interactions.

### A. ACTR as a sequence-specific self-avoiding random walk

By employing only bond and excluded volume interactions, model 1 treats ACTR as a self-avoiding polymer, similar to standard bead-spring polymer models. The main difference here is that the excluded volume interactions are highly specific (represented at a united-atom level of resolution), such that they induce some amount of sequence specificity into the model. Figure 2(a) shows the distribution of *R*_g_ values for ACTR generated by simulations of model 1 (blue curve) at a reduced temperature 0.87*T*^∗^. *T*^∗^ is defined as the temperature at which the model reproduces 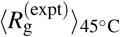. For model 1, ⟨*R*_g_⟩ is approximately independent of temperature, as expected for a self-avoiding random walk under athermal solvent conditions^12^. For this reason, and since there is no free interaction parameter in model 1 for reproducing 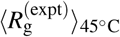, we cannot directly define *T*^∗^ in this case. However, the value of the temperature-independent ⟨*R*_g_⟩ for model 1 is 26.4 *Å* (dashed blue line in Figure 2(a)), which is nearly the same as the experimentally measured 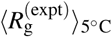. Therefore, we can interpret this model as representing an ensemble at 0.87*T*^∗^ ([5°C + 273°C]*/*[45°C + 273°C] ≃0.87).

**FIG. 2:**
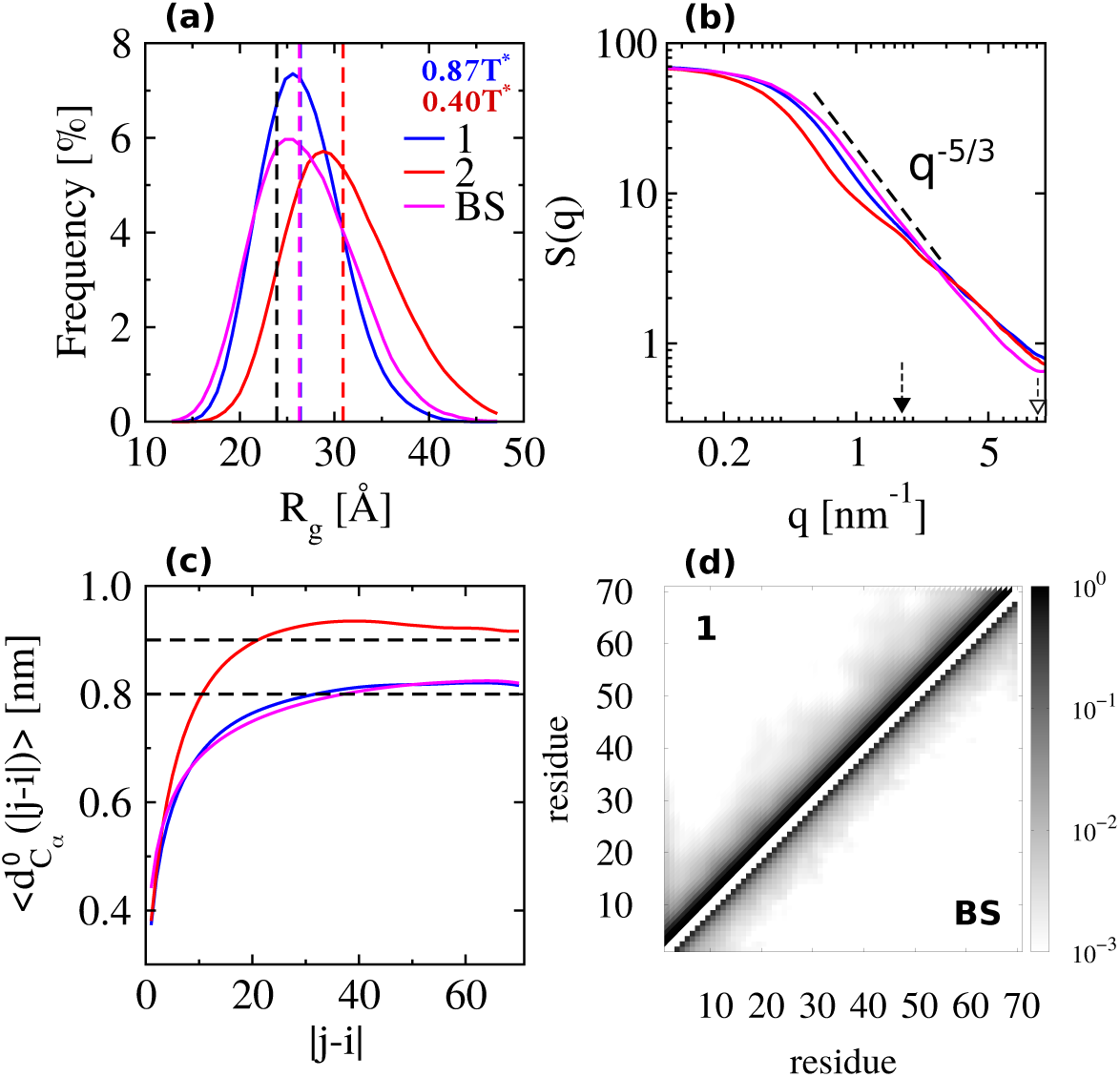
(a) Distribution of the radius of gyration, *R*_g_, (b) single-chain backbone structure factor, *S*(*q*), (c) root mean square normalized distance between pairs of residues separated by | *j* −*i*| residues along the chain, 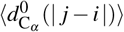, and (d) the probability of pairs of *C*_*α*_ atoms to be within a cutoff of 1.0 nm. In panel (a) the dashed black line indicates the experimental result of ⟨*R*_g_ ⟩ at 45°C. In panels (a)-(c), the blue, red and magenta curves correspond to results from model 1, model 2 and the BS model, respectively. The arrows in panel (b) indicate the value of *q* at which the scaling law of *S*(*q*) changes for model 1 (filled arrow) and for the BS model (empty arrow). In panel (d), the top and bottom triangles correspond to results from model 1 and the BS model, respectively.

Panels (b)-(d) of Figure 2 present various ensemble-averaged properties of model 1 at 0.87*T*^∗^ (blue curves). Note that the average fraction of helical segments per residue is negligible in this model, due to the lack of interactions that stabilize helices (see Figure S6). Panel (b) presents the structure factor, *S*(*q*), which describes the overall shape of the protein at three characteristic length scales^64^. For small 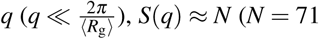 for ACTR). For 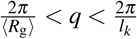, a power law of *S*(*q*) ∼ *q*^−1*/v*^ occurs, where *v* describes the quality of the solvent according to standard polymer theory. The so-called Kuhn length, *l*_*k*_, is model dependent. For 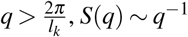 corresponding to a rigid rod. For model 1, *l*_*k*_ ≈ 2.2 nm, since the crossover to rigid rod scaling occurs at approximately *q* ∼ 2.9 nm^−1^ (filled arrow in Figure 2(b)). Additionally, *ν* ≈ 3*/*5 in region 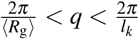, indicating that the conformational ensemble generated by model 1 is comparable to a polymer in good solvent (i.e., extended conformations are prominent). Panel (c) presents the root mean square (normalized) distance between C_*α*_ atoms for two residues separated by |*j* − *i*| residues along the chain, 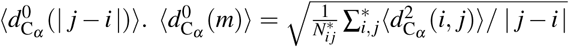, where 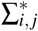 is a sum over all *i j* pairs with | *j* − *i* |= *m*, 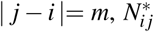 is the number of such pairs, and 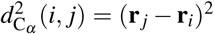. Note the normalization by | *j* − *i* |, in contrast to other, related work^77^ (see Methods section for further details and Figures S7 and S8 for plots of the unnormalized root mean square distances and additional analysis of 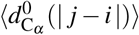, respectively). 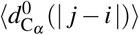 characterizes the local concentration of peptide segments for short separation distances 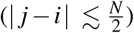 and the global behavior of the chain for larger separation distances (|*j* − *i*| ∼ *N*). For model 1, 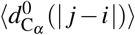 increases monotonically as a function of |*j* − *i*|, reaching a value of approximately 0.82 nm at |*j* − *i*| = *N*. This behavior is very similar to that of a self-avoiding random walk (magenta curve, discussed further below), and thus consistent with the analysis of *S*(*q*). Panel (d) of Figure 2 presents the probability that a particular pair of residues, *i* and *j*, are in contact (i.e., their *C*_*α*_ atoms are within 1 nm of one another). The top left triangle of the plot corresponds to the conformational ensemble generated by model 1, displaying a very low probability of two residues being in contact if they are situated more than a few residues from one another along the chain. In other words, the chain is very extended, in further support of the results from the shape parameters.

To compare more directly with a standard polymer model, we also simulated a bead-spring (BS) polymer model, commonly referred to as the Kremer-Grest model^61^. Figure S9 demonstrates the temperature-independent distribution of *R*_g_ values for this model. We aligned the length scale of the models by applying a rescaling factor (0.45 nm) to the BS model such that 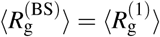. Note that the temperature of the BS model (*T*^∗^ = 2.0 *ε/k*_*B*_) was chosen such that the width of the distribution of *R*_g_ values approximately reproduced that of model 1. In the BS model, residues interact according to a purely repulsive (i.e., WCA) potential and the bonds between neighboring beads are represented with a FENE potential. Thus, the main difference between the models is the accuracy with which model 1 describes the excluded volume of both the backbone and the side chains. Figure 2 demonstrates that, with the exception of a broader distribution of *R*_g_ values (panel (a)), shorter Kuhn length (*l*_*k*_ ≈ 0.66 nm, indicated by the empty arrow in panel (b)), and a modest change in the probabilities of contact for neighboring residues (panel (d)), the conformational ensemble of the BS model is very similar to the ensemble generated by model 1, according to these metrics. We also compared the gyration tensors from the BS model and from model 1. The gyration tensor eigenvalues and normalized asphericity values, 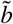, are given in Table II. As shown in Figure 3, the ratio of the gyration tensor eigenvalues is *λ*_3_ : *λ*_2_ : *λ*_1_ = 12.20 : 3.13 : 1 for the BS model compared with *λ*_3_ : *λ*_2_ : *λ*_1_ = 11.81 : 3.12 : 1 for model 1, further confirming the self-avoiding random walk behavior generated by model 1. Additionally, the ensembles generated by these models yield similar asphericity values: 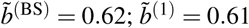.

**TABLE II:** Eigenvalues of the gyration tensor and normalized asphericity values.

| model id | $\lambda_3$ [nm<sup>2</sup>] | $\lambda_2$ [nm<sup>2</sup>] | $\lambda_1$ [nm<sup>2</sup>] | $\bar{b}$ |
| --- | --- | --- | --- | --- |
| BS | 4.88 | 1.25 | 0.40 | 0.62 |
| 1 | 5.08 | 1.34 | 0.43 | 0.61 |
| 2 | 6.47 | 1.76 | 0.55 | 0.61 |
| BS-LJ | 3.58 | 1.41 | 0.57 | 0.47 |
| 3a | 3.54 | 1.20 | 0.41 | 0.53 |
| 3b | 3.80 | 1.19 | 0.42 | 0.55 |
| 4 | 3.56 | 1.14 | 0.39 | 0.55 |
| 5a | 3.55 | 1.15 | 0.40 | 0.54 |
| 5b | 3.51 | 1.12 | 0.39 | 0.55 |
| 6 | 3.38 | 1.04 | 0.36 | 0.56 |

**FIG. 3:**
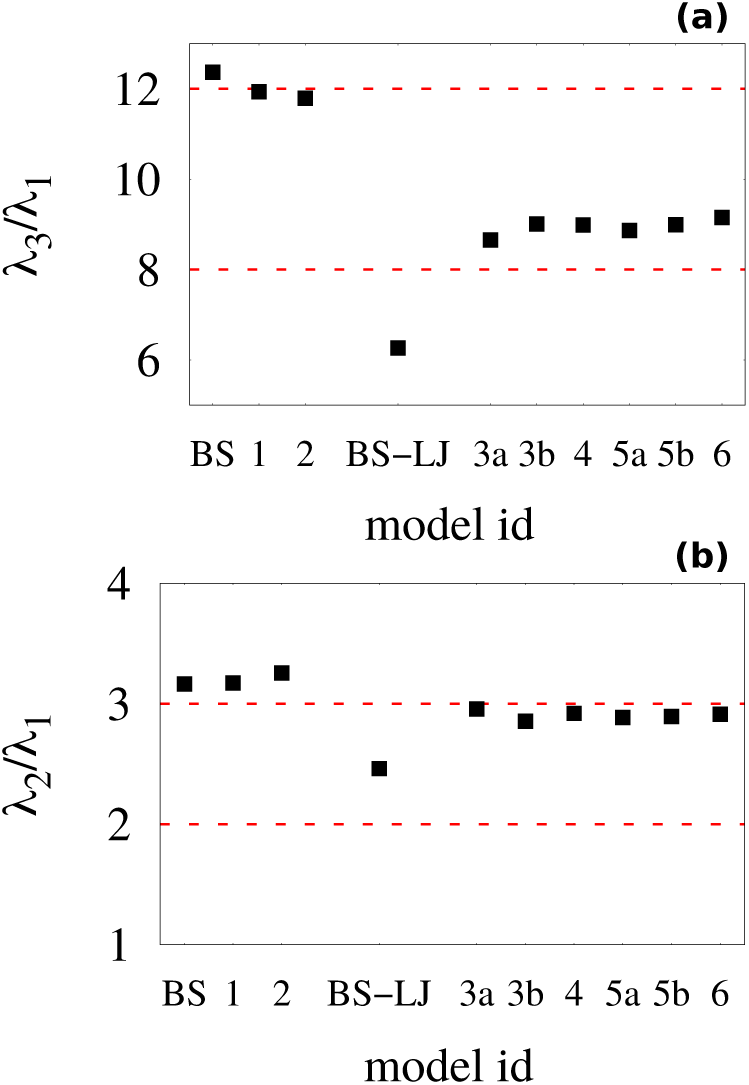
The ratio of eigenvalues of the gyration tensor: (a) *λ*_3_*/λ*_1_; (b) *λ*_2_*/λ*_1_.

Figure 2 also presents properties generated from simulations of model 2 (red curves). In contrast to model 1, model 2 introduces an effective backbone stiffness into the set of interactions which results in an overall expansion of the peptide for comparable absolute temperatures. In fact, for this particular model, 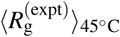 is too low to reproduce at any temperature due to the fixed nature of the effective stiffness of the backbone, as determined by the Amber99SB-ILDN force field. Nevertheless, to illustrate the overall properties of the model, Figure 2 presents results from 0.4*T*^∗^, with 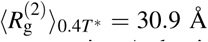 (dashed red line in panel (a)). In this case, *T*^∗^ was approximated via a linear extrapolation of ln *R*_g_(*T*) (i.e., assuming Arrhenius behavior). Panel (b) demonstrates that model 2 has similar properties to model 1, i.e., the peptide behaves approximately as a polymer in good solvent. However, the crossover to *S*(*q*) ∼ *q*^−1^ occurs at a smaller *q* compared with model 1, indicating that the addition of backbone stiffness results in a larger approximate *l*_*k*_, as expected. The contact probability maps of the two models are also quite similar (Figure S10). However, panel (c) of Figure 2 demonstrates more clearly the effect of local backbone stiffness. In particular, 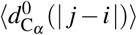 grows more quickly for | *j* −*i*| ≤ 40, compared with model 1, and then drops slowly to a value of about 0.92 nm at | *j* −*i*| = *N*. The peak at | *j* −*i*| ≈ 40 indicates that the chain is locally more rigid in model 2, while the larger distance at | *j* −*i*| = *N* is indicative of more extended conformations overall, as seen in panel (a). We also compared the gyration tensor for these models (Figure 3). The ratio of the gyration tensor eigenvalues for model 2 are *λ*_3_ : *λ*_2_ : *λ*_1_ = 11.76 : 3.20 : 1, again demonstrating similar behavior to model 1. The ensemble generated by model 2 also has comparable asphericity to the ensemble generated by model 1 (see Table II).

To obtain a more detailed picture of the conformational landscapes of these models, we performed a dimensionality reduction using the UMAP nonlinear manifold learning algorithm^71^ to determine a two-dimensional embedding upon which to view the ensembles. UMAP attempts to retain both the local pairwise connectivity as well as the overall global structure of the high-dimensional input space, within a lower-dimensional (e.g., two-dimensional) projection. As input features for this procedure, we employed distances between pairs of segments along the peptide and angles between triplets of segments, as described in the Methods section. For consistent comparisons, the two-dimensional UMAP embedding was determined from simulations of model 4, and subsequently applied to the other conformational ensembles. Rows (a) and (b) of Figure 4 demonstrate an approximate physical interpretation of each of the embedding dimensions. Panel (bi) presents a scatter plot of points sampled along the embedding, with colors corresponding to the *R*_g_ of each conformation. There is a significant correlation between UMAP-1 and *R*_g_, although this relationship is notably nonlinear. Additionally, the distribution of conformations is significantly broader along UMAP-1 compared with *R*_g_. The second dimension is more difficult to directly interpret. Panel (bii) presents a scatter plot with colors corresponding to the average angle formed between the first, seventh, and eleventh segments, when the peptide is partitioned into segments of four residues. Row (a) presents an illustration of this angle for two representative conformations. As one moves from the lower left to the upper right of the embedding space, conformations display an overall transition from extended structures to more hairpin like structures. The UMAP landscape provides a clearer view of the heterogeneous ensemble of structures sampled by ACTR, compared with, e.g., free-energy landscapes plotted as a function of *R*_g_ and *R*_e_ (Figure S11). The nonlinear nature of this embedding results in structured free-energy landscapes, which are often not possible for disordered ensembles using linear techniques^69,70^. Row (c) of Figure 4 presents the free-energy landscapes along the embedding for model 1 and for the BS model. Both models appear to sample very similar conformational ensembles (of primarily extended, larger *R*_g_, structures), consistent with the analysis of the shape parameters above.

**FIG. 4:**
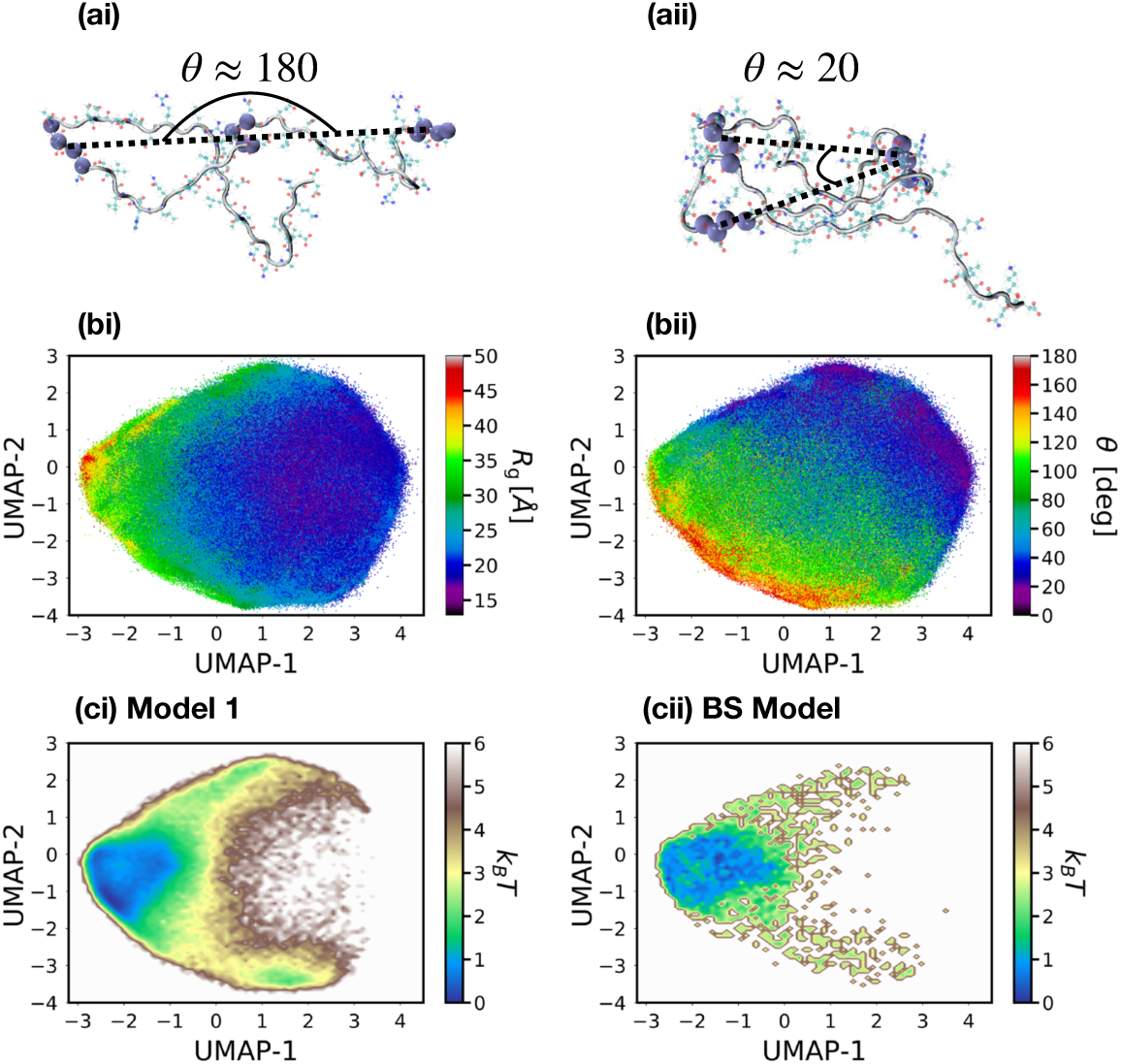
(a) Illustrations of the angle, *θ*, formed between the first, seventh, and eleventh segments, when the peptide is partitioned into segments of four consecutive residues along the backbone. (b) Heat maps of (i) *R*_g_ and (ii) *θ* along the coordinates determined from the UMAP manifold learning algorithm. (c) Free-energy landscapes generated by (i) model 1 and (ii) the BS model along the UMAP coordinates.

### B. The effect of hydrophobic attraction between side-chains

Models 3a, 3b, and 4 go beyond the simple self-avoiding walk picture by incorporating attractive interactions between *C*_*β*_ atoms to represent the solvent-induced hydrophobic attraction between amino acid side chains. While models 3b and 4 take into account the relative hydrophobicity of each residue and scale this hydrophobic attraction accordingly, model 3a employs a uniform hydrophobic attraction which reproduces the average hydrophobicity of the peptide chain. In addition to hydrophobic attractions, model 4 incorporates explicit electrostatic interactions between charged residues via the Debye-Hückel formalism. Figure 5 presents a comparison of the properties generated by these models at *T*^∗^. Panel (a) presents the distribution of *R*_g_ values for models 3a, 3b, and 4 as the blue, red, and orange curves, respectively. The distributions are nearly identical, although model 4 has a slight tendency towards more collapsed structures. This demonstrates an insensitivity in the overall dimensions of the peptide to changes in specific interactions between residues (given the constraints enforced by the excluded volume interactions). Similar to models 1 and 2, the formation of helices is negligible for these models (Figure S6). However, these models no longer demonstrate properties of a polymer in good solvent (panels (b) and (c)). In particular, *S*(*q*) displays *ν* = 1*/*2 dependence, representing a polymer in Θ solvent. In other words, the attractive hydrophobic interactions approximately counteract the effect of excluded volume and chain stiffness, resulting in random walk behavior.

**FIG. 5:**
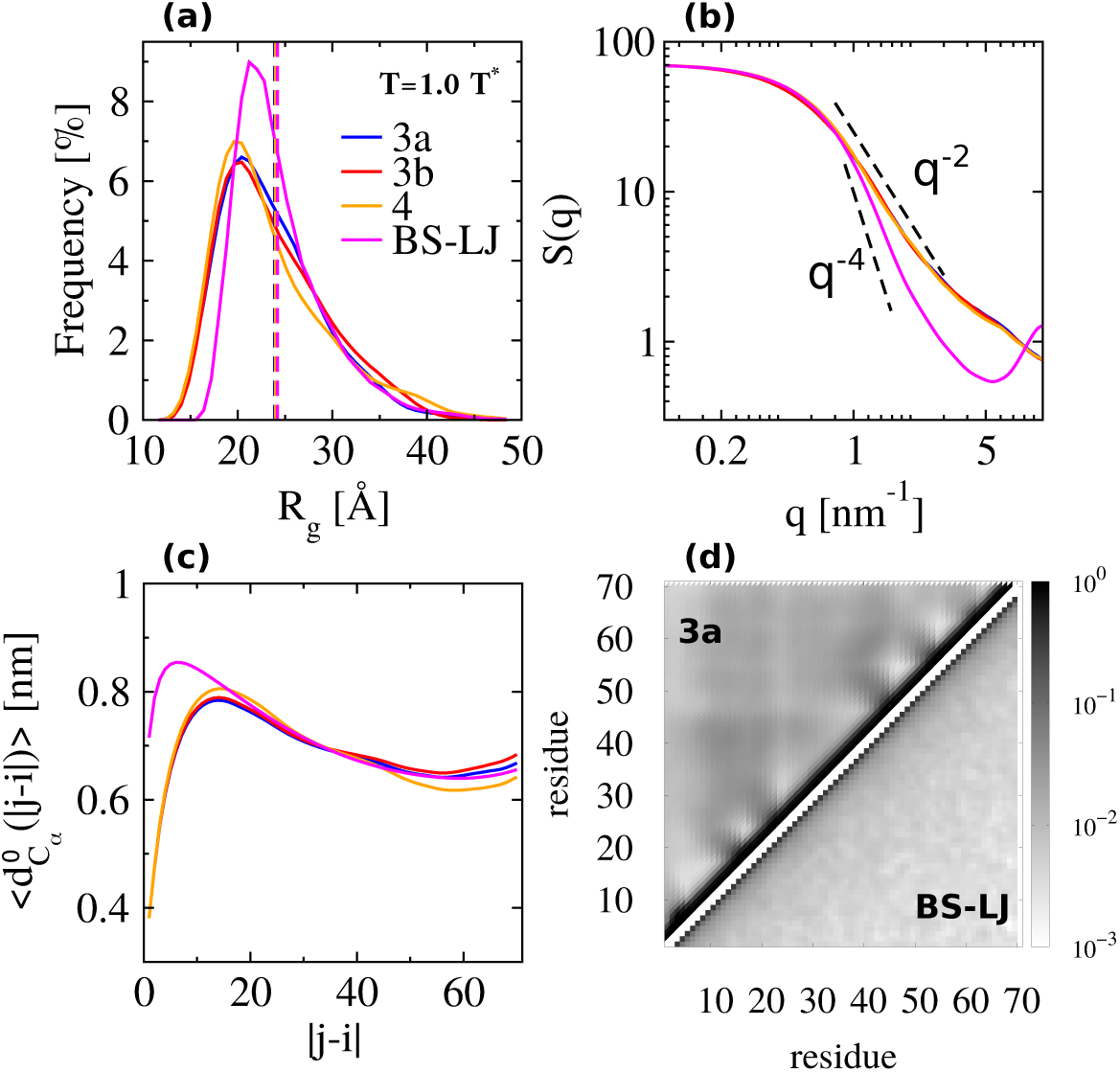
(a) Distribution of the radius of gyration, *R*_g_, (b) single-chain backbone structure factor, *S*(*q*), (c) root mean square normalized distance between pairs of residues separated by | *j* −*i*| residues along the chain, 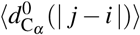, and (d) the probability of pairs of *C*_*α*_ atoms to be within a cutoff of 1.0 nm. In panel (a) the dashed black line indicates the experimental result of ⟨*R*_g_ ⟩ at 45°C. In panels (a)-(c), blue, red, orange and magenta curves correspond to results from model 3a, model 3b, model 4 and the BS-LJ model, respectively. In panel (d), the top and bottom triangles correspond to results from model 3a and the BS-LJ model, respectively.

Panel (c) also demonstrates notable differences of these conformational ensembles, relative to the self-avoiding random walks. In particular, 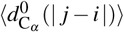 displays a maximum at | *j* −*i*|≈ 15, which reflects the local rigidity of the chain due to the backbone stiffness (as seen for model 2). As | *j*−*i*| increases beyond 15, 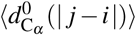 decreases until a minimum is reached at | *j*−*i*|≈ 55, due to the attractive hydrophobic interactions between side chains which promote more collapsed structures. Finally, the slight increase of 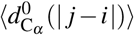 for larger | *j* − *i* | values demonstrates persistent conformational heterogeneity (i.e., the ensemble is not completely collapsed). Models 3a and 3b demonstrate very similar behavior, although a slight expansion of distances is observed in model 3b over the entire range of | *j* −*i*| separations. The inclusion of electrostatics in model 4 results in noticeable compaction of the ensemble for larger | *j* −*i*| separations. This result may seem surprising, since ACTR has a − 8 net charge. However, recall that we have calibrated the energy scale of each model by adjusting the absolute simulation temperature to match *R*_g_ with the experimental value. In this case, the direct effect of adding electrostatics to the model does indeed result in a shift in the *R*_g_ distribution to larger values if the absolute simulation temperature remains fixed, as expected from the net charge on the chain. By considering the models at *T*^∗^, we demonstrate that *given ensembles with fixed* ⟨*R*_g_⟩, the ensemble generated by the model with electrostatics samples somewhat more compact structures.

We again compare these ensembles with a standard polymer model (BS-LJ), but incorporate attractive interactions between monomers, as described in the Methods Section. The obtained distribution of *R*_g_ values as a function of temperature can be seen in Figure S9. We again aligned the length scale of the models by applying a rescaling factor (0.73 nm) to the BS-LJ model such that 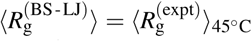. The distribution of *R*_*g*_ generated by the BS-LJ model is presented in panel (a) of Figure 5 (magenta curve), showing a narrower distribution and fewer very compact structures compared with model 3a. This may be partially due to the fact that we have not re-optimized the temperature for the BS-LJ model (*T*^∗^ = 2.0 *ε/k*_*B*_) to fit the width of the distribution of *R*_g_ values. Significant differences are also observed in *S*(*q*) (panel (b) of Figure 5), which demonstrates *v* = 1*/*4 behavior, indicating that the chain behaves more like a polymer under poor solvent conditions in the BS-LJ model (i.e., samples overall more compact conformations). This result is consistent with previous work with this model, which identified the Theta temperature as approximately *T*^∗^ = 3.0 *ε/k*_*B*_^78^. *The S*(*q*) behavior appears to be in conflict with the distribution of *Rg* (panel (a) of Figure 5), which is narrower than the distribution generated by model 3a, without sampling the compact tail of the distribution from model 3a. However, panel (c) of Figure 5 demonstrates that although a maximum in 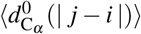 occurs at short residue separations in the BS-LJ model, due to a lack of interactions governing local stiffness of the chain, larger 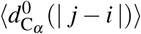 values are also attained in this region. These larger average distances between residues at short separation along the chain likely prevent the sampling of structures with the smallest *R*_g_ values. At the same time, the lack of chain rigidity along with the presence of attractive interactions between monomers together promote an increased sampling of compact structures, leading to apparently compact behavior at intermediate length scales. Additional distinctions between the two ensembles can be seen by examining the ratios of the gyration tensor eigenvalues, which are *λ*_3_ : *λ*_2_ : *λ*_1_ = 6.28 : 2.47 : 1 for the BS-LJ model and *λ*_3_ : *λ*_2_ : *λ*_1_ = 8.63 : 2.93 : 1 for model 3a (Figure 3). Moreover, the ensemble generated by the BS-LJ model 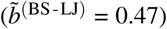 is slightly more spherical than the ensemble generated by model 3a 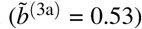. Figure 5(d) presents the contact probability maps generated by model 3a (upper left) and the BS-LJ model (lower right). While both models display increased probability of long-separation (along the chain) contacts, relative to the models without attractive interactions, the comparison highlights the simplicity of the BS-LJ ensemble relative to the ensemble generated by model 3a. In contrast to the slightly more expanded ensembles generated by models 1 and 2, sequence-specific excluded volume interactions (along with the details of local protein chain stiffness) appear to play a more significant role in determining the finer details of these more collapsed conformational ensembles. On the other hand, the contact probability maps of models 3a, 3b, and 4 display relatively smaller deviations from one another (Figure S10). Overall, the inclusion of attractive interactions results in a structured contact probability map, but remains largely independent of the precise distribution of hydrophobic attractions.

Column (i) of Figure 6 presents the free-energy landscapes for models 3a, 3b and 4, plotted along the UMAP embedding introduced above. The most striking difference between these landscapes compared to those generated by the self-avoiding walk models is the expanded diversity of structures sampled despite rather similar distributions of *R*_g_. The addition of attractive interactions results in sampling both more collapsed and more expanded structures compared with model 1. There are also more subtle differences between the conformational ensembles generated by models 3a, 3b, and 4. The redistribution of hydrophobicity in model 3b, compared with model 3a, leads to only a slight shift in the conformational ensemble, as indicated by the analysis above. The most prominent difference is perhaps the increased sampling of the smallest UMAP-1 (largest *R*_g_) values, although this difference manifests itself as only a minor change in the overall distribution of *R*_g_. There is also an increase of structures corresponding to the largest values of UMAP-1. Overall, the differences between the ensembles generated by models 3a and 3b appear to be distributed throughout the entire embedding space, resulting in “averaging out” and little difference in the overarching features of the disordered ensembles. However, the introduction of electrostatics (model 4) leads to more significant differences in the ensemble of structures and, in particular, a more rugged free-energy landscape (i.e., a larger number of clearly separated local minima), as seen in panel (ci) of Figure 6. ACTR has 18 charged residues, 5 of them are positive charges and 13 are negatively charged (see Equation (1)). Overall, the electrostatic interactions lead to increased sampling of compact structures (positive values of UMAP-1), and also a slight increase in structures with the smallest UMAP-1 (largest *R*_g_) values. The conformations along UMAP-2 (i.e., with different *θ* values) appear to more uniformly affected by the addition of electrostatic interactions. It should be noted that the calibration of the energy scales through the use of reduced temperatures, as discussed above, results in a distinct balance of stiffness versus attractive interactions in the different models. In the case of model 4 (and for model 6 below), a lower absolute simulation temperature is required for this model to reproduce the appropriate ⟨*R*_g_⟩ value, resulting in larger stiffness energies relative to *k*_B_*T*^∗^. This difference in absolute simulation temperatures might be interpreted as the reason for the larger difference in the free-energy landscape for model 4, compared with models 3a and 3b. Alternatively, one can say that given the fixed model details (e.g., chain stiffness, hydrophobicity, etc.), the ensemble which incorporates electrostatics and reproduces 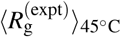 does so through an increase in the ensemble ruggedness.

**FIG. 6:**
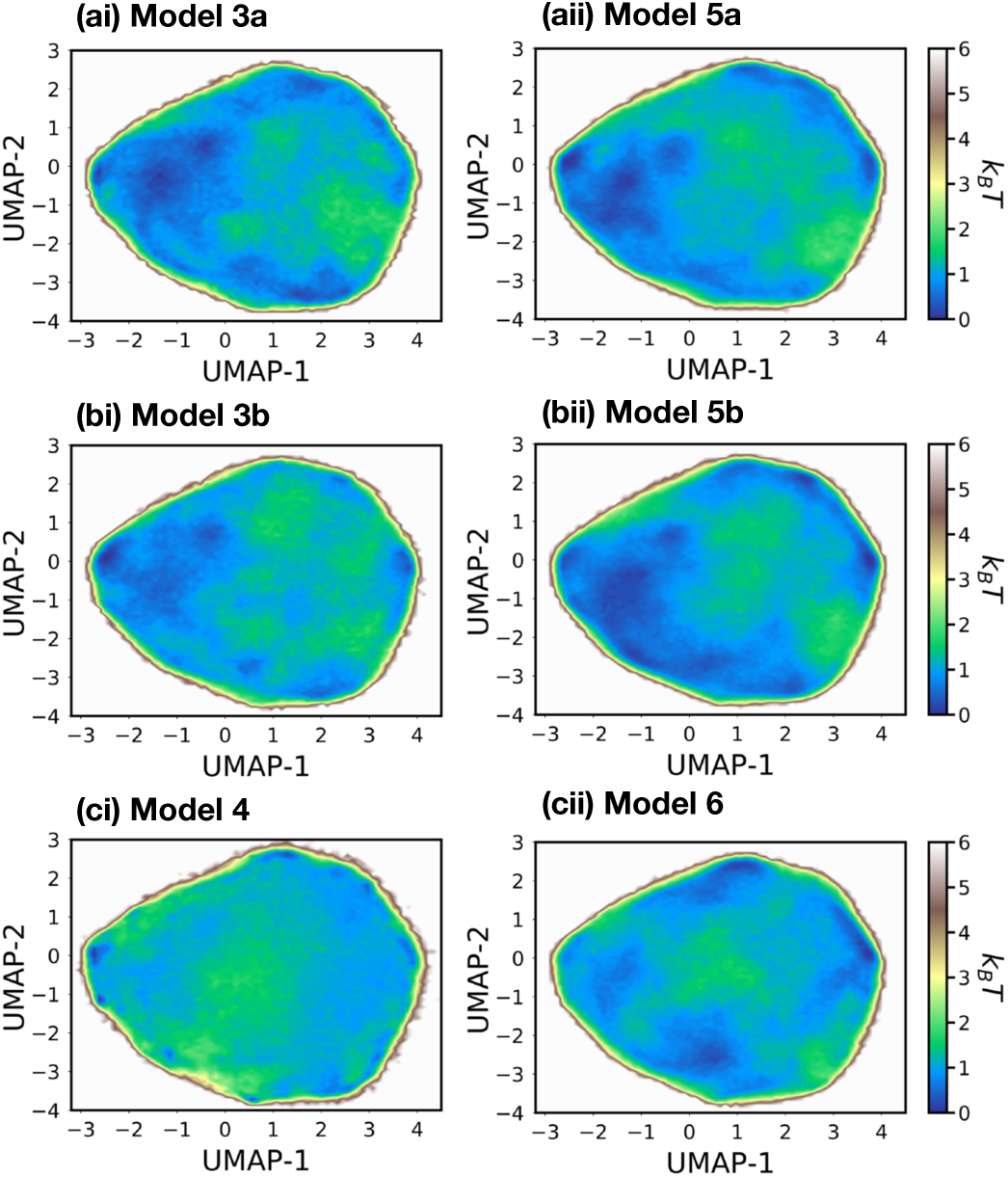
Free-energy landscapes generated by models 3a, 3b, and 4 (column i) and models 5a, 5b, and 6 (column ii) along the coordinates determined from the UMAP manifold learning algorithm.

### C. Transient helices

In addition to hydrophobic attractions between side chains, models 5a, 5b and 6 employ attractive interactions between *C*_*α*_ atoms separated by three peptide bonds along the protein backbone in order to represent hydrogen-bonding interactions. The parameter for this interaction was chosen to approximately reproduce the overall propensity for helices in ACTR, as measured in experiments (described further in the Methods Section). The current models do not include hydrogen-bonding-like interactions between residues farther apart along the peptide chain, since propensity toward *β* -sheet-like secondary structures have not been observed in ACTR. Similar to the previous set of models, model 5a employs uniform hydrophobic interactions, while models 5b and 6 use residue-specific hydrophobicity parameters. Additionally, model 6 incorporates electrostatic interactions between charged residues. Figure 7(a) presents the distribution of *R*_g_ values at *T*^∗^ for models 5a, 5b, and 6 as the red, blue, and orange curves, respectively. We find that the distributions are rather insensitive to the addition of hydrogen-bonding interactions. These models generate SAXS profiles and corresponding Kratky plots in good agreement with experimental measurements (see Figure S12 compared with Figure 2b of^7^).

**FIG. 7:**
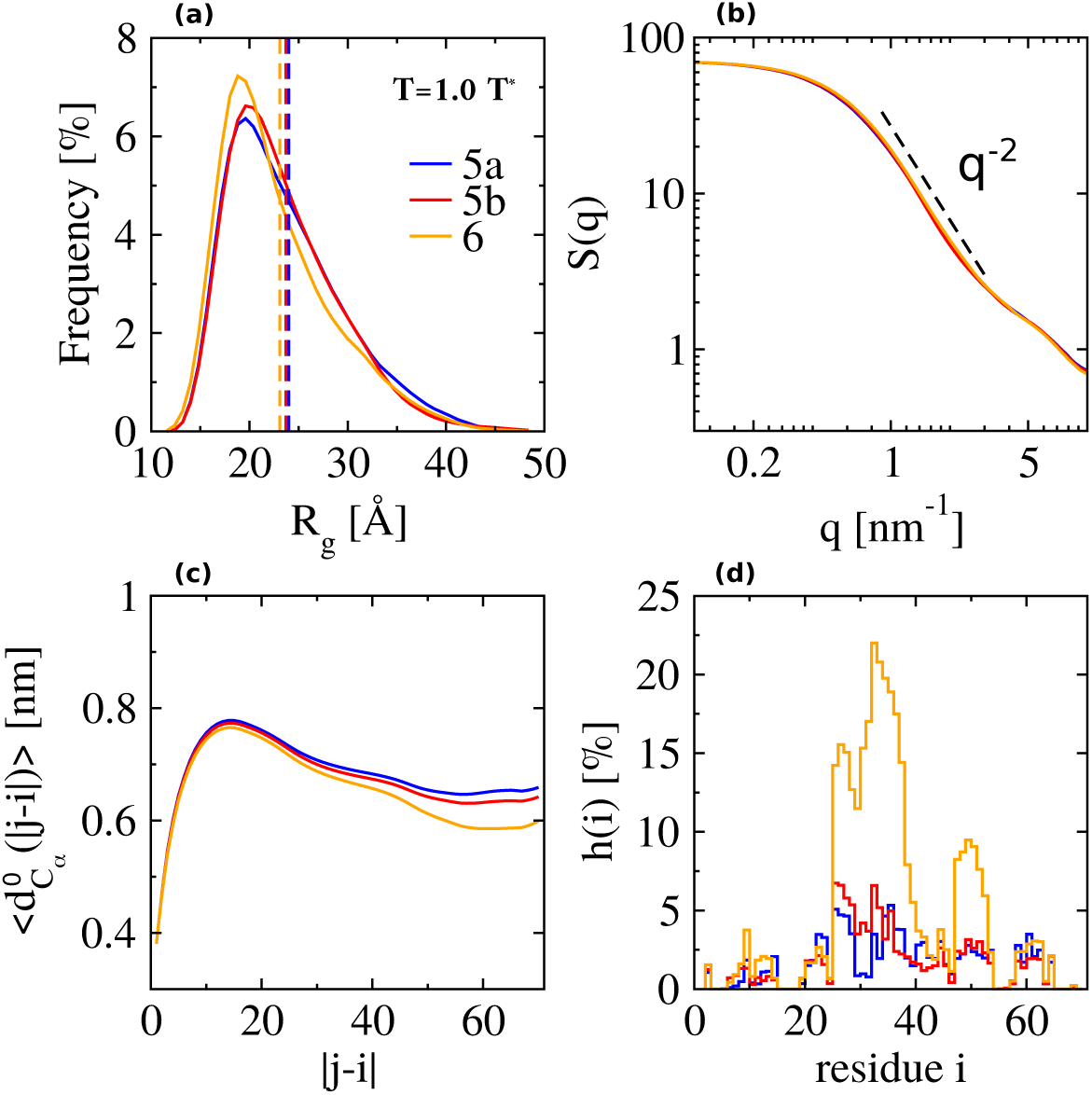
(a) Distribution of the radius of gyration, *R*_g_, (b) single-chain backbone structure factor, *S*(*q*), (c) root mean square normalized distance between pairs of residues separated by | *j* − *i* | residues along the chain, 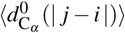, and (d) the average fraction of helical segments, *h*(*i*). In panel (a) the dashed black line indicates the experimental result of ⟨*R*_g_ ⟩ at 45°C. In panels (a)-(d), blue, red and orange curves correspond to results from model 5a, model 5b and model 6, respectively.

Panels (b) and (c) of Figure 7 present *S*(*q*) and 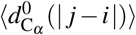, respectively, for models 5a, 5b and 6. No significant differences are observed in the behavior of *S*(*q*), which can be fit to *q*^−2^, i.e., a polymer in Θ solvent. Panel (c) demonstrates that the behavior of 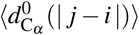 is insensitive to the inclusion of hydrogen-bonding interactions, i.e., 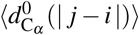 follows the same trend as for models 3a, 3b and 4. However, similar to the case of model 4, 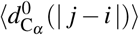 for model 6, which includes electrostatics, is smaller than for models 5a and 5b for all separation distances | *j* −*i*|. Additional differences in the ensembles generated by these three models can be seen by examining the gyration tensor. As shown in Figure 3, the ratio of the gyration tensor eigenvalues is *λ*_3_ : *λ*_2_ : *λ*_1_ = 8.87 : 2.87 : 1 for model 5a, 9.00 : 2.87 : 1 for model 5b, and 9.39 : 2.89 : 1 for model 6. Overall, these results indicate that incorporating hydrogen-bonding interactions causes a slight shift in the ensembles towards self-avoiding walk behavior, although the conformational ensemble as a whole still behaves like a random walk (per *S*(*q*)). Additionally, the addition of electrostatics amplifies this effect through increased stabilization of helices, as examined in more detail below. At the same time, the ensembles remain largely spherical (see Table II). The contact probability maps for these models are presented in Figure S10, but exhibit differences similar to those between the models without hydrogen-bonding interactions. We characterize the formation of helices by the propensity of each residue to form a helical segment, *h*(*i*), as described in the Methods section. Figure 7(d) presents *h*(*i*) for models 5a, 5b and 6 (blue, red, and orange curves, respectively). All three models demonstrate similar behavior in terms of the position of helix formation along the chain, due to the accurate treatment of side chain excluded volume. For example, there is dip in the region with residue indices from 28 to 30, likely due to the presence of arginine with residue index 29, which is hydrophilic and contains a rather bulky side chain. The helical regions at residue positions [9:14], [29:40], [48:54] and [58:62] are in agreement with experimental observations^6,79^. The helical content of models 5a and 5b appears to be somewhat insensitive to the distribution of hydrophobic interactions, indicating that the precise hydrophobic contacts play a limited role in the formation of helices (given the fixed representation of sterics). On the other hand, model 6 demonstrates significantly larger helicity. This may be due to either (i) the generic stabilization of helices from the increased compaction of the ensemble or (ii) the increased contact of specific residues which then promote the formation of helices, which we discuss further below.

The free-energy landscapes for models 5a, 5b and 6, plotted along the UMAP embedding introduced above, are presented in Figure 6. Similar to the results for 3a and 3b, the UMAP projections for models 5a and 5b are quite comparable. In contrast to the previous set of models, while model 6 does slightly focus the sampling toward particular regions of the landscape, the ensemble does not appear as rugged as for model 4. However, a clear view of the ensemble is perhaps clouded by the helical conformations, since the UMAP coordinates were determined based on an ensemble without helical conformations. To obtain a more detailed picture of the formation of helices, we performed a dimensionality reduction using principal component analysis (PCA) while employing the backbone dihedral angles as input features (i.e., dihedral PCA^72^). Although the ensembles are largely disordered, linear dimensionality reduction can effectively characterize the formation of transient helices within these ensembles. Panels (b)-(d) of Figure 8 present the free-energy surfaces generated by models 5a, 5b, and 6, respectively, along the first two PCs. A clustering was performed along the first three PCs, in order to partition the conformational space into 50 microstates. Here, we present a coarse-grained view of this clustering, attained by grouping together sets of microstates. The coarse cluster definitions are presented in Figure 8(a) as a function of the first two PCs. Figure 9 characterizes each cluster with the intra-cluster *h*(*i*) distributions. Cluster A (blue curve) represents structures with a small helix formed from residues 25-40, while cluster E (gray curve) contains structures with negligible helical conformations. There are two pathways from the A cluster to the E cluster which sample either negative (a) or positive (b) values of PC-2. Figure 9 demonstrates that pathway (a) corresponds to unraveling the helix from the N-terminus (panel (a)), while pathway (b) corresponds to unraveling the helix from the C-terminus (panel (b)). The free-energy surfaces in Figure 8 show that models 5a and 5b sample a single dominant pathway for helix formation, while the introduction of electrostatics in model 6 allows for helix formation from either end. This additional pathway leads to a significant increase in the sampling of helical conformations (Figure 7(d)).

**FIG. 8:**
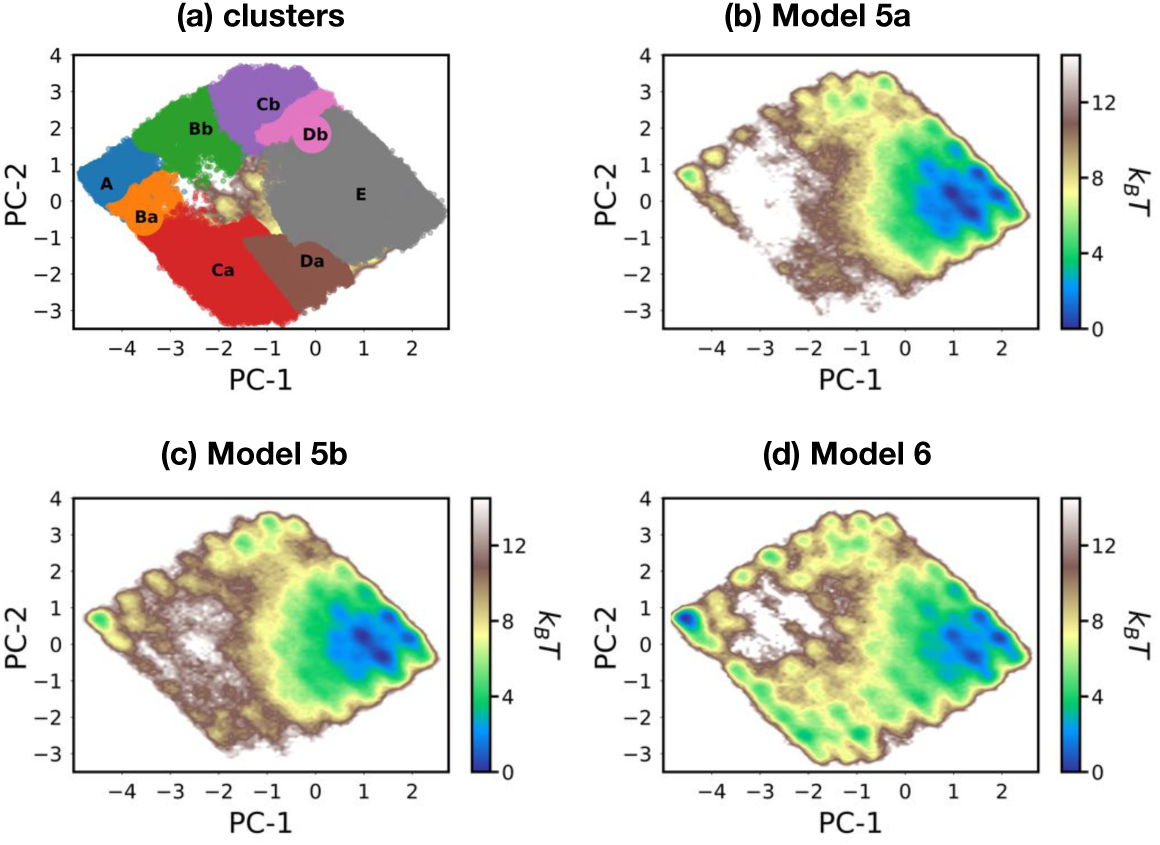
(a) Conformational clusters of ACTR presented along the two dominant principal components (PCs). (b)-(d) The free-energy surfaces of ACTR generated using models 5a, 5b and 6, plotted along the two dominant PCs.

**FIG. 9:**
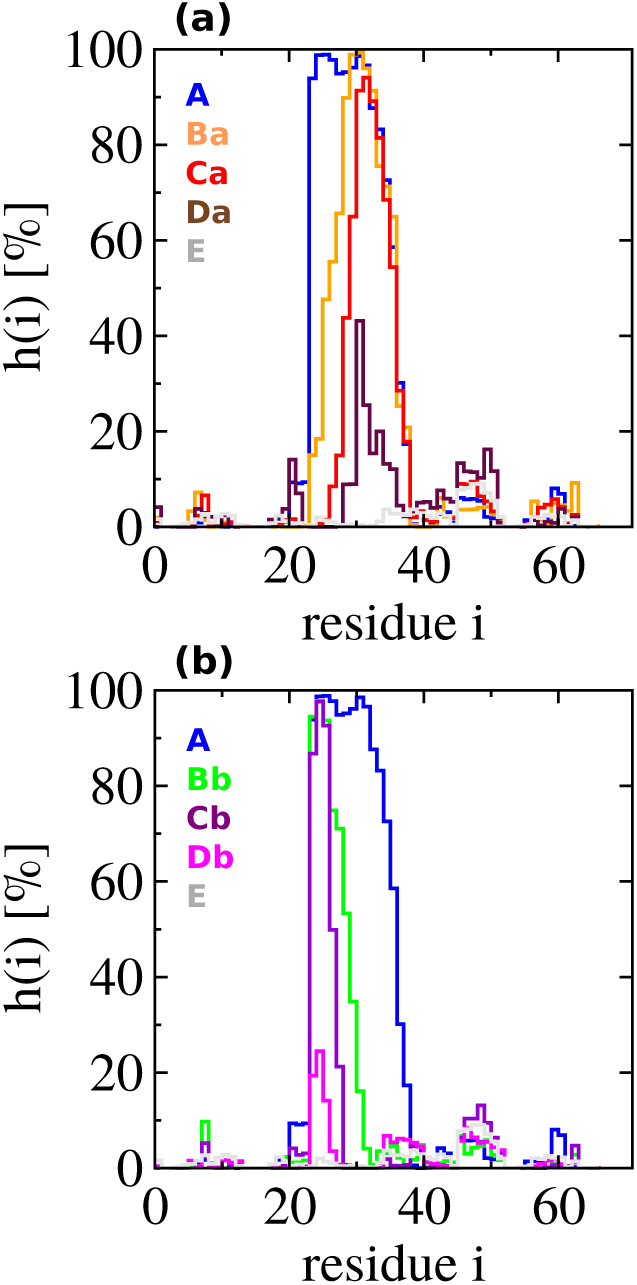
Intra-cluster fraction of helical segments, *h*(*i*), for (a) N-terminus and (b) C-terminus folding pathways, as characterized on the PCA landscape (Figure 8) using model 6.

### D. Clarifying the role of excluded volume in the formation of helical structures

We have considered the impact that both generic and specific attractive interactions have on the resulting conformational ensembles of peptides with the length and approximate excluded volume of ACTR. To explicitly demonstrate the role that the steric interactions play, we consider models for the uncharged polypeptides (Alanine)_71_ and (Glycine)_71_, denoted as polyA and polyG, respectively, which have the same parameters *ε*_hp_ and *ε*_hb_ as model 5a, but lack the sequence-specific side-chain sterics of ACTR. Because the resulting ensembles are dramatically different from one another, it is not feasible to match 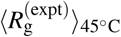 by adjusting the temperature, as performed for the other models in this study. Instead, we employ the same absolute simulation temperature as for model 5a, allowing *R*_g_ to deviate from the experimental value.

Figure 10 presents the average fraction of helical segments, *h*(*i*), for ACTR, polyA, and polyG. In going from the ACTR to the polyA model, all side-chain atoms except the C_*β*_ atoms are removed. The removal of excluded volume interactions reduces the entropy loss upon helix formation, significantly promoting the sampling of helical conformations. By further removing the C_*β*_ atoms, from polyA to polyG, the attractive interactions which stabilize compact structures (including helices) are removed, resulting in a complete absence of helical conformations. Despite the uniformity of the sequence, the helicity of polyA demonstrates sequence-dependent behavior. The two regions of smaller helicity at [25 : 30] and [47 : 52] (black lines in Figure 10) arise due to the likelihood of the chain bending at positions corresponding to 1/3 and 2/3 of the total chain length, in order to maximize hydrophobic contact in compact structures (see, e.g., Figure S13). Thus, even if one were to reparametrize the model to reproduce the appropriate ⟨*R*_g_⟩ value and overall *h*(*i*) magnitude, the formation of helices in the models that do not accurately represent the side chain sterics would be qualitatively incorrect. Furthermore, in contrast to the small differences in the overall “shape” of the protein for the various models with differing energetics considered above, there is a dramatic change in the conformational ensemble associated with the amendment of excluded volume interactions (Figure 10(b)). This motivates the use of models that accurately represent the protein sterics for the investigation of disordered conformational ensembles.

**FIG. 10:**
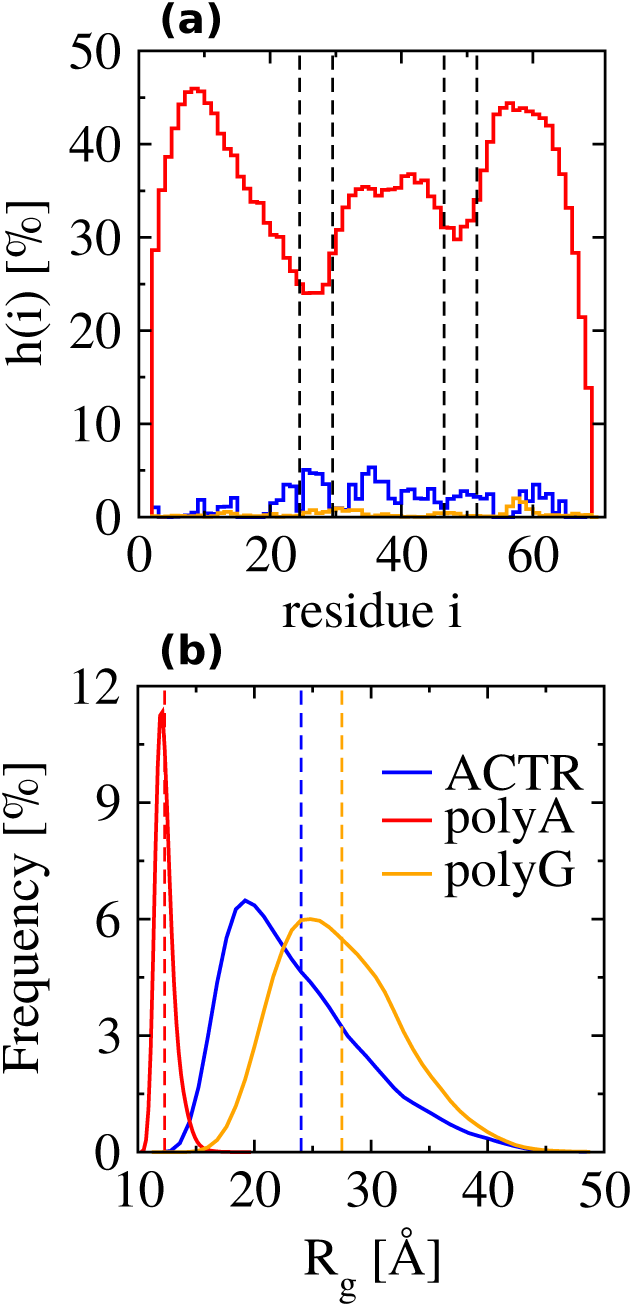
(a) Average fraction of helical segments, *h*(*i*), and (b) distribution of the radius of gyration, *R*_g_, for ACTR (blue), polyA (red) and polyG (orange), determined from simulations of model 5a. The two regions marked by the black lines in panel (a) include the residues ranges [25 : 30] and [47 : 52].

### E. The conformational ensemble of NCBD

To investigate the applicability of the the considered models for investigating distinct disordered ensembles, we consider NCBD—the binding partner of ACTR. NCBD has 59 residues, with 27 hydrophobic and 8 charged residues (see Equation (2)). Its unbound conformer (PDB-id: 2KKJ) forms a molten globule that has three helices at residue positions [6 : 19], [23 : 36], and [36 : 47] (see Figure 1 for helix positions in the bound state).^35,50^ Thus, the unbound NCBD protein generates a very distinct conformational ensemble compared with ACTR. In fact, NCBD and ACTR are representative examples of two different classes of IDPs^77^ (see Figure S2(b)). The ⟨*R*_g_⟩ for NCBD was measured from SAXS experiments to be approximately 18.8 Å under native-like conditions.^6^ We consider here only models 5b and 6, to investigate whether electrostatics play a significant role in shaping the unbound conformational ensemble of NCBD. Because initial simulations of these models resulted in a lack of helix stabilization, we increased the energy of the hydrogen-bonding-like interaction, *ε*_hb_, from 13 to 16.9 (30% larger than that of ACTR), which lead to good agreement of both ⟨*R*_g_⟩ and *h*(*i*) with respect to the experimental values. We have again calibrated the energy scale of the model by finding the simulation temperature at which the experimental values of ⟨*R*_g_⟩ and *h*(*i*) are reproduced, independently from ACTR, although the resulting *T*^∗^ is only 10% larger (in absolute temperature units, i.e., K) than the value for ACTR. The adjustment of parameters to reproduce the properties of NCBD was expected, since these quantities are free-energy functions which rigorously depend on the system identity and thermodynamic state point^80^. In fact, the relative insensitivity of the model parameters indicates a certain level of transferability of the model, further motivating the use of simple energetic functions for representing disordered ensembles.

Figure 11(a) presents the distribution of *R*_g_ for models 5b (red curve) and 6 (orange curve), which are nearly identical (*R*_g_ = 18.7 and 18.3 *Å*, respectively). Similarly, *S*(*q*) (panel (b)) demonstrates *v* = 1/2 behavior for both models, indicating that electrostatics play a relatively small role in the overall shape of NCBD. However, Figure 11(c) demonstrates that 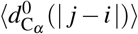 is significantly different for models 5b and 6 for | *j* −*i*| > 15, qualitatively similar to the comparison of models 5b and 6 for ACTR. Without electrostatics (model 5b), NCBD demonstrates less compaction in the intermediate regime due to the onset of attractive interactions, compared with ACTR. The inclusion of electrostatics (model 6) leads to a greater degree of compaction for NCBD in this regime, and a significant difference in 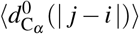 generated by the two models. The eventual increase of 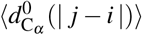 demonstrates that NCBD retains significant conformational heterogeneity within its molten globule ensemble, despite the presence of largely formed helices. The difference in the behavior of 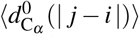 for the two models in the case of NCBD is striking, considering the similarity of the ensemble in terms of ⟨*R*_g_⟩, *S*(*q*), and *h*(*i*). This may be a result of the slightly lower propensity for middle helices in model 6 (panel (d)), which can allow for the sampling of more compact structures through stacking of the outer helices. However, the gyration tensor provides further evidence of the similarity of the ensembles generated by models 5b and 6. The ratio of the gyration tensor eigenvalues is 9.75 : 3.35 : 1 and 9.78 : 3.04 : 1 for models 5b and 6, respectively, while the normalized asphericity values are 0.53 and 0.56. Overall, it appears that the conformational ensembles of IDPs with large fractions of secondary structure motifs may be more robust to perturbations in the interactions, assuming a fixed representation of sterics.

**FIG. 11:**
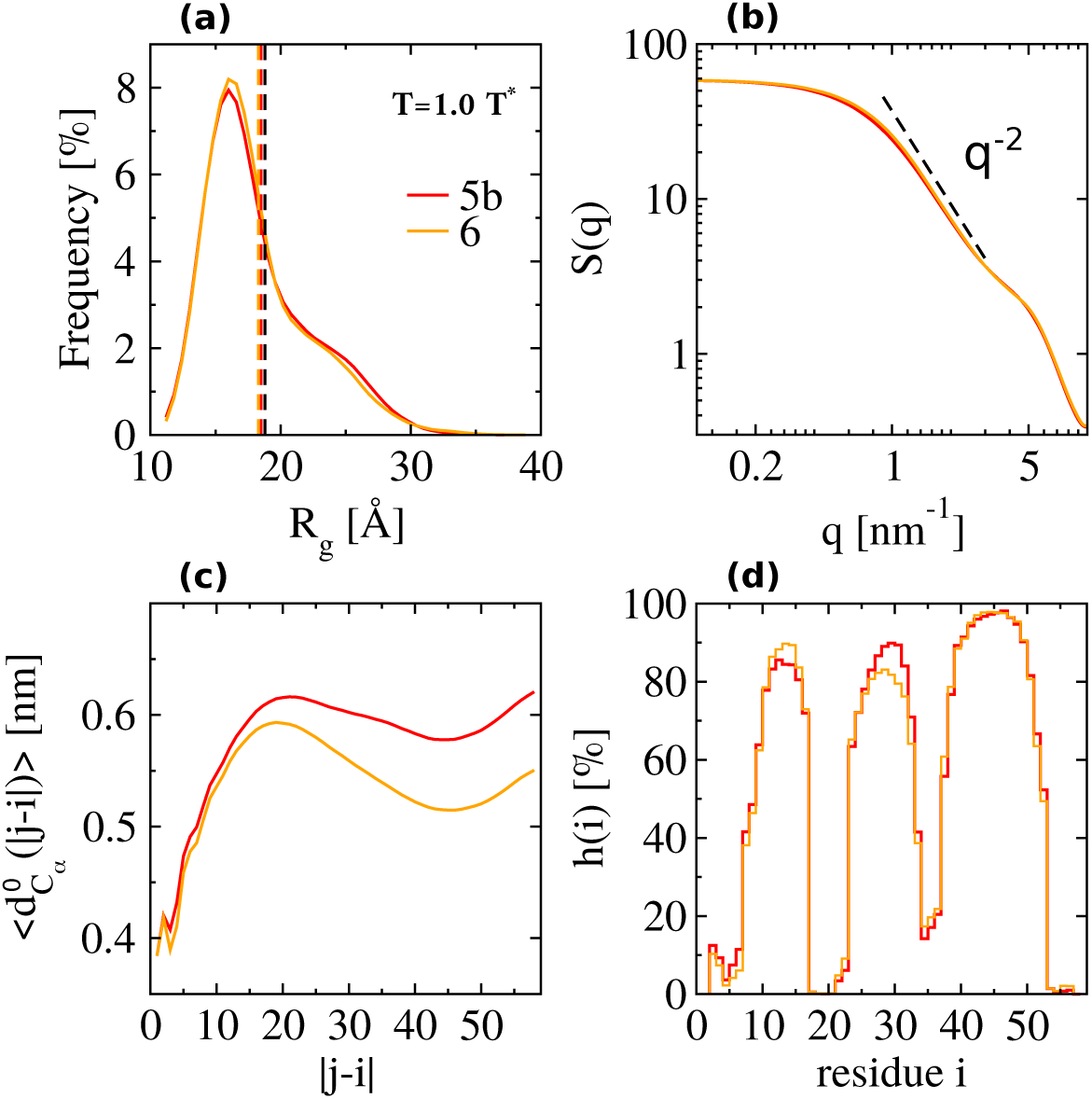
(a) Distribution of the radius of gyration, *R*_g_, (b) single-chain backbone structure factor, *S*(*q*), (c) root mean square normalized distance between pairs of residues separated by | *j* − *i* | residues along the chain, 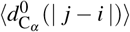, and (d) the average fraction of helical segments, *h*(*i*). In panel (a) the dashed black line indicates the experimental result of ⟨*R*_g_ ⟩. In panels (a)-(d), red and orange curves correspond to results from model 5b and model 6, respectively.

## IV. Conclusions

We have studied the ensembles of two intrinsically disordered peptides, ACTR and NCBD, using a simple physics-based model, which accurately represents peptide sterics and allows an adjustable parametrization to match experimental quantities, e.g., ⟨*R*_g_⟩ and *h*(*i*). A hierarchy of models was considered, which systematically incorporated an increasing number and complexity of interactions, in order to clarify the impact of these interactions on the features of the resulting ensembles. Our analysis demonstrates that the differences between these distinct ensembles are difficult to fully characterize using only traditional shape parameters, such as the distribution of radius of gyration values and the single-chain backbone structure factor. However, the root mean square normalized inter-residue distances between C_*α*_ atoms, the ratio of gyration tensor eigenvalues, and the contact probability map assist in further distinguishing the overarching features of the ensembles. Additionally, we have employed a manifold learning algorithm in this work, to determine an optimal two-dimensional representation for viewing the ensemble of conformations, which provides an effective way to further clarify the differences between distinct disordered ensembles.

Our investigation found that, with respect to a self-avoiding random walk, disordered ensembles that incorporate hydrophobic interactions lead to a significant increase in conformational heterogeneity. However, given the presence of attractive interactions, the precise identity of these interactions, e.g., the distribution of hydrophobic interactions along the chain or the presence of electrostatics, appear to play a relatively small role in determining the major features of the disordered free-energy landscapes. At the same time, specific interactions can stabilize particular structures which may be relevant for processes under a perturbation of the system (e.g., when a disordered peptide comes into contact with its binding partner). For example, electrostatic interactions increase the ruggedness of the free-energy landscape and stabilize multiple routes to secondary structure formation. These effects appear to be more significant for more disordered, flexible IDPs (e.g., ACTR), than for molten globules (e.g., NCBD). While electrostatics are thought to play an important role in the formation of encounter complexes in IDPs^38,39^, the present work suggests that specific contacts between charged residues can promote the presence of transient helices within the ensemble of conformations sampled in solution, which may be relevant for coupled folding and binding processes.

The flexible physics-based model employed in this work facilitated the reproduction of experimental ⟨*R*_g_⟩ and *h*(*i*) values for both ACTR and NCBD. These two peptides are representative examples of two different classes of IDPs: “fully disordered” (ACTR) and molten globule (NCBD). Although the (free-energy) parameters of this simple model should be, in principle, highly sequence specific, we find that only relatively small adjustments were necessary to reproduce the experimental measurements for both systems. This indicates a certain level of transferability in terms of the essential features shaping the free-energy landscape for these disordered systems, motivating the continued use of coarse-grained models. Moreover, in conjunction with previous investigations of helix-coil transitions^33,34^, our results indicate that excluded volume interactions play a key role in determining the overarching characteristics of heterogeneous landscapes. This further motivates the development of models that can accurately model protein sterics while efficiently sampling conformational space.

## Supporting Information Available

The Supporting Information provides additional model and simulation details as well as further analysis.

## Acknowledgments

The authors thank Hsiao-Ping Hsu and Govardhan Reddy for critical reading of the manuscript. Y.Z. and J.F.R. thank Tristan Bereau and Hsiao-Ping Hsu for fruitful discussions. J.F.R. thanks Yasemin Bozkurt Varolgüneş for assistance with the UMAP calculations. J.F.R is very grateful to Ben Schuler and his group for insightful discussions regarding the NCBD/ACTR system. This work was partially supported by European Research Council under the European Union’s Seventh Framework Programme (FP7/2007-2013)/ERC Grant Agreement No. 340906-MOLPROCOMP, and by the Deutsche Forschungsgemeinschaft (DFG, German Research Foundation) - Project number 233630050 - TRR 146.

