## Supporting Information for "Investigating the Conformational Ensembles of Intrinsically-Disordered Proteins with a Simple Physics-Based Model"

### Model Details

#### Bonded interactions given by the Amber99 SB-ILDN force field

The potential for two covalently bonded atoms $i$ and $j$ with distance $r_{ij}$ is represented by a harmonic potential $V_b(r_{ij}) = \frac{1}{2}k^b_{ij}(r_{ij} - b_{ij})^2$, where $b_{ij}$ is the equilibrium distance and $k^b_{ij}$ is the harmonic force constant. The angle potential between three continuous atoms $i$, $j$ and $k$ is represented by a harmonic potential $V_a(\theta_{ijk}) = \frac{1}{2}k^\theta_{ijk}(\theta_{ijk} - \theta^0_{ijk})^2$, where $\theta^0_{ijk}$ is the equilibrium angle and $k^\theta_{ijk}$ is the force constant. The potential for dihedral angles $\phi$ between two planes $ijk$ and $jkl$ composed by four continuous atoms $i$, $j$, $k$ and $l$ is defined as $V_d(\phi_{ijkl}) = k_\phi(1 + \cos(n\phi - \phi_s))$, where $\phi_s$ is the equilibrium dihedral angle and $k_\phi$ is the force constant. The parameters $k^b_{ij}$, $b_{ij}$, $k^\theta_{ijk}$, $\theta^0_{ijk}$, $k_\phi$, multiplicity $n$ and $\phi_s$ are directly taking from the Amber99SB-ILDN force field without modification. Example plots of $V_b(r_{ij})$, $V_a(\theta_{ijk})$ and $V_d(\phi_{ijkl})$ can be found in Fig. S1.

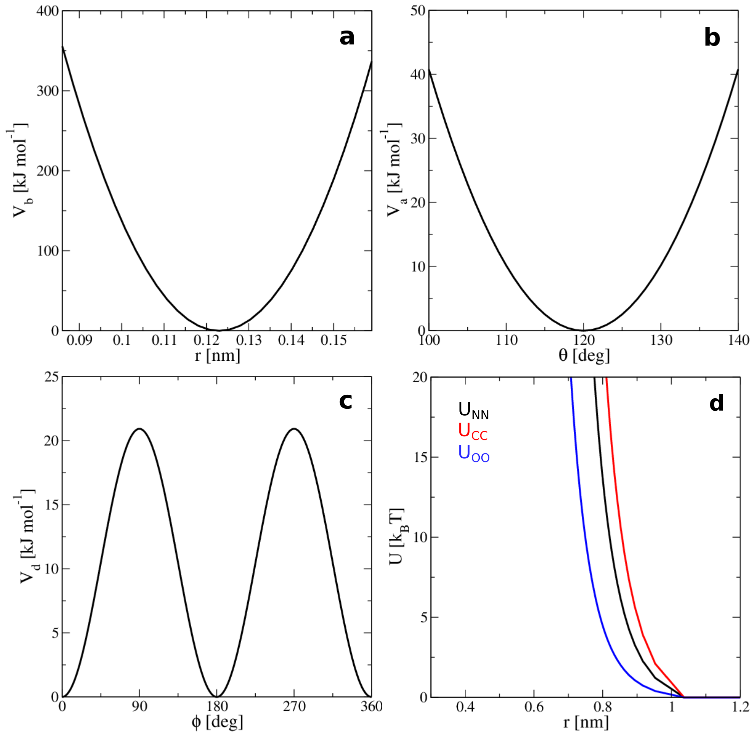


Figure S1: Functional forms for the bond (a), angle (b), and dihedral (c) potentials. Panel (d) presents the WCA potentials employed to model nitrogen-nitrogen (NN), carbon-carbon (CC) and oxygen-oxygen (OO) interactions.

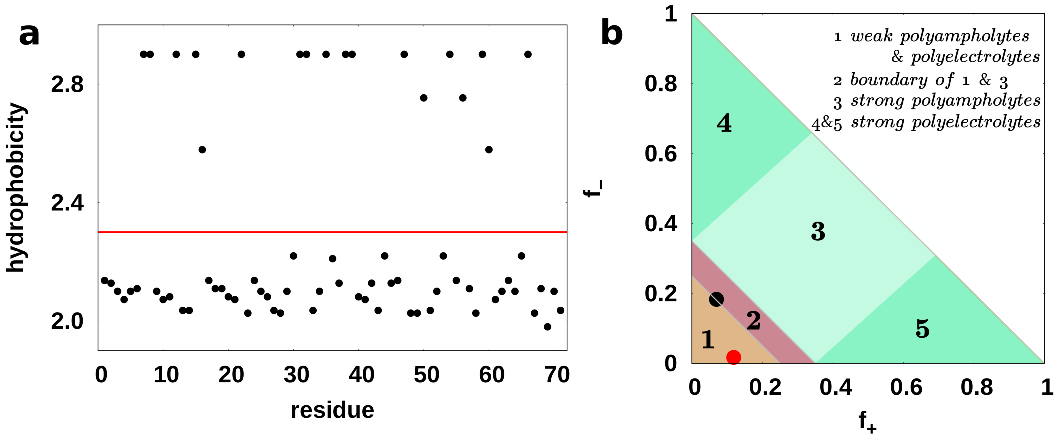


Figure S2: (a) The hydrophobicity scale (i.e., $\epsilon_{\mathrm{hp},i}$ values for each residue $i$) for the ACTR sequence, employed for the models with hetero-hydrophobic interactions. The red line denotes $\epsilon_{\mathrm{hp}} = 2.3$, used for the models with homo-hydrophobic interactions. (b) IDP classification diagram recreated from Fig. 7 in[1] . ACTR and NCBD are represented by the black and red circles, respectively.

#### Excluded volume effects and 1-4 interactions

In the physics-based CG model considered in this work, the excluded volume interactions of (heavy) atoms along the peptide backbone are represented with Weeks-Chandler-Andersen (WCA) potentials (Fig. S1(d)), constructed directly from the Lennard-Jones interactions of the AMBER99SB-ILDN force field. Exclusions were treated per the AMBER99SB-ILDN force field, i.e., $i/i+1$ and $i/i+2$ pairs were excluded, while the interaction for $i/i+3$ (i.e., 1-4) pairs were reduced by a factor of 0.5 (i.e., fudgeLJ = 0.5 in the terminology of the GROMACS simulation suite).

#### Hydrophobicity scale and IDP classification of ACTR

The models employing hetero-hydrophobic interactions scale the attraction between $C_\beta$ atoms according to the residue identities. The hydrophobicity scale, based on the Miyazawa-Jernigan interaction matrix,[2] is shown in Fig. S2(a) and in Table S1. The magnitude of the scale was calibrated such that the average hydrophobicity (see the red line in panel (a)) is identical to that of the corresponding homo-hydrophobic interaction. Panel (b) of Fig. S2 presents a classification diagram for IDPs, recreated from Fig. 7 in[1]. Here, IDPs are grouped according to their net charge per residue (NCPR), NCPR $= \mid f_+ - f_- \mid$, where $f_+$ and $f_-$ are the fractions of positive and negative charges. NCBD has an NCPR of 0.44, falling into region 1, which consists of weak polyampholytes and weak polyelectrolytes that often exhibit globule-like ensembles. ACTR has an NCPR of 0.75, falling into region 2—an boundary region between region 1 and the region consisting of strong polyampholytes. Region 2 contains IDPs that tend to sample heterogeneous conformational ensembles, including many of the known proteins which undergo couple folding and binding.

**Table S1: The hydrophibic energies of residues used in the physics-based model.**

| residue | $\epsilon_{\mathrm{hp},i}$ | residue | $\epsilon_{\mathrm{hp},i}$ |
|---|---|---|---|
| LYS | 1.981 | HIS | 2.211 |
| GLU | 2.027 | ALA | 2.220 |
| ASP | 2.036 | TYR | 2.431 |
| ASN | 2.073 | CYS | 2.477 |
| SER | 2.082 | TRP | 2.569 |
| ARG | 2.100 | VAL | 2.579 |
| GLN | 2.100 | MET | 2.597 |
| PRO | 2.109 | ILE | 2.753 |
| THR | 2.128 | PHE | 2.873 |
| GLY | 2.137 | LEU | 2.901 |

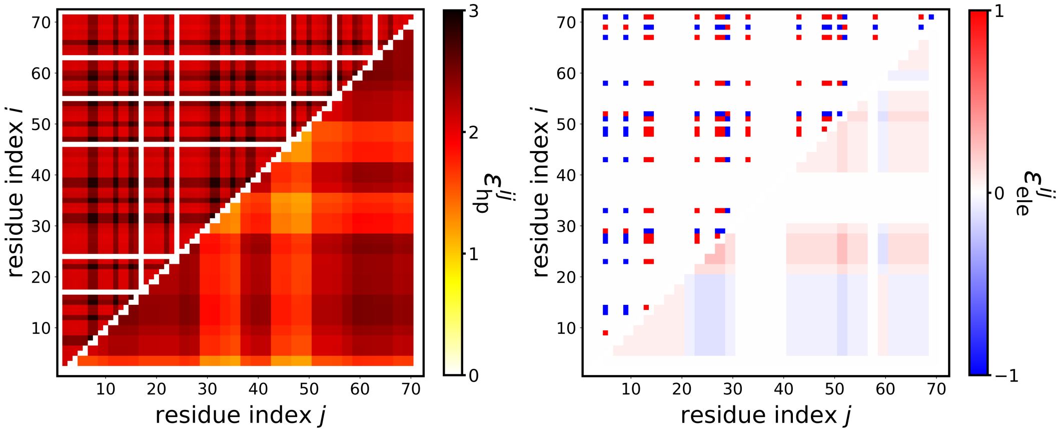


Figure S3: The pairwise interaction strength for hydrophobic (left) and electrostatic (right) interactions. The top and bottom triangles correspond to the "full" coarse-grained resolution and a coarser resolution obtained by partitioning the peptide chain into segments of four resides and averaging over pairs belonging to these segments, respectively.

#### Pairwise hydrophobic and electrostatic interactions

Fig. S3 presents the pairwise interaction energy for the hydrophobic and electrostatic interactions. The blank regions in panel (a) indicate the location of glycine residues, which do not have $C_\beta$ atoms, thus lacking hydrophobic attractions. The upper triangles show the interaction strengths at the residue-level of resolution, while the lower triangles present a coarse-grained view, obtained by partitioning the peptide chain into segments of four resides and averaging over pairs belonging to these segments.

### Simulation Convergence

#### Autocorrelation functions

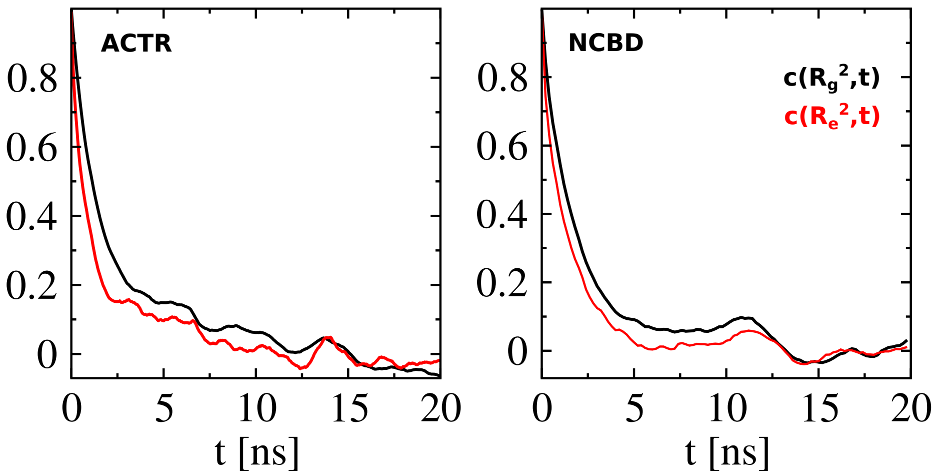


Figure S4: The autocorrelation function of $R_\mathrm{g}^2$ and $R_\mathrm{e}^2$ for ACTR (left) and NCBD (right) determined from simulations of model 5b at $T^*$.

We test the convergence of the trajectories by calculating the autocorrelation function $c(A,t) = \frac{\langle A(t_0)A(t_0+t)\rangle - \langle A(t_0)\rangle\langle A(t_0+t)\rangle}{\langle A(t_0)^2\rangle - \langle A(t_0)\rangle^2}$, where $t$ is the lag time and $A$ is an observable. Fig. S4 presents the autocorrelation functions of the mean square gyration radius $c(R_\mathrm{g}^2, t)$ and mean square end-to-end distance $c(R_\mathrm{e}^2, t)$ for ACTR and NCBD.

#### Sub-trajectories

We also randomly divided each trajectory into two groups and calculated $R_g$ and $h(i)$ for each subtrajectory (Fig. S5).

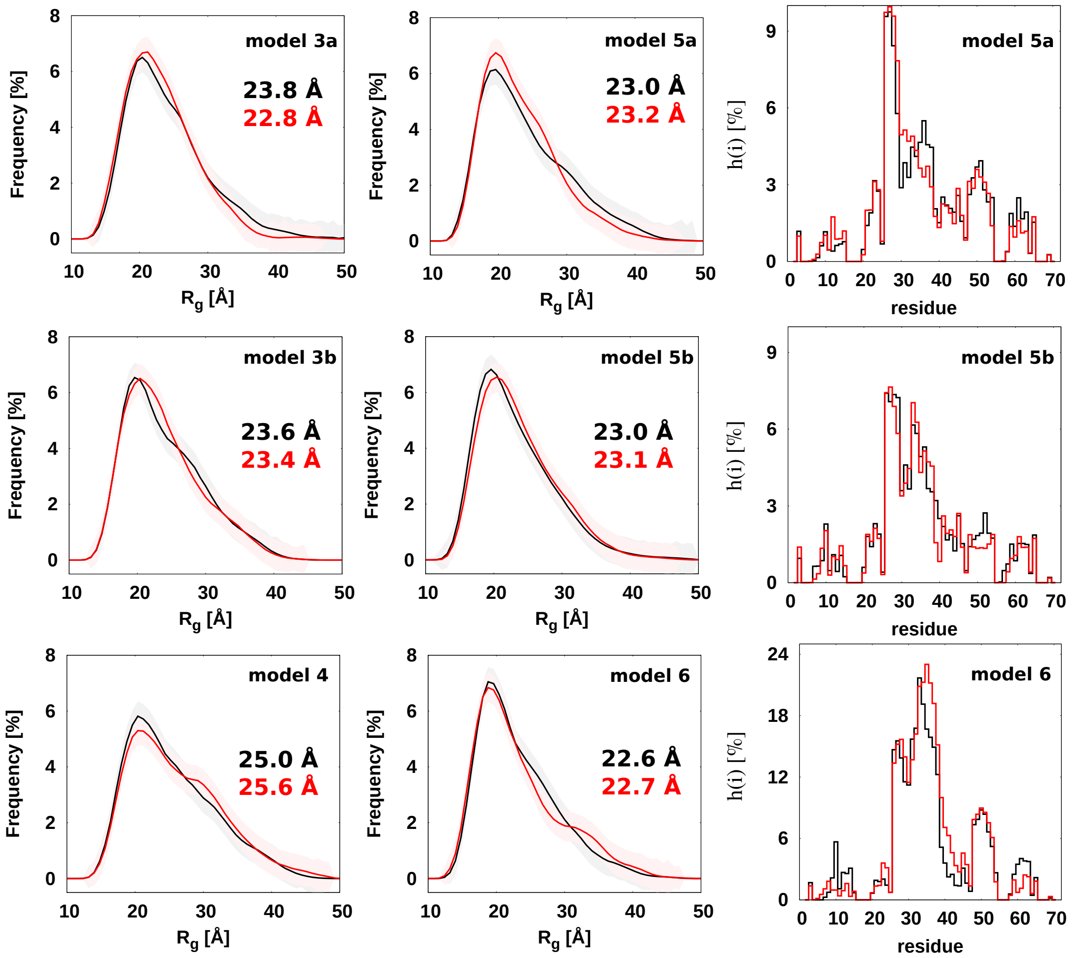


Figure S5: The trajectories of ACTR are divided into two groups to check the convergence of the simulations, the calculated $R_g$ and $h(i)$ from the two groups are denoted by the black and red colors, respectively.

### Additional Results

#### Helicity of ACTR in models without hydrogen-bonding-like interactions

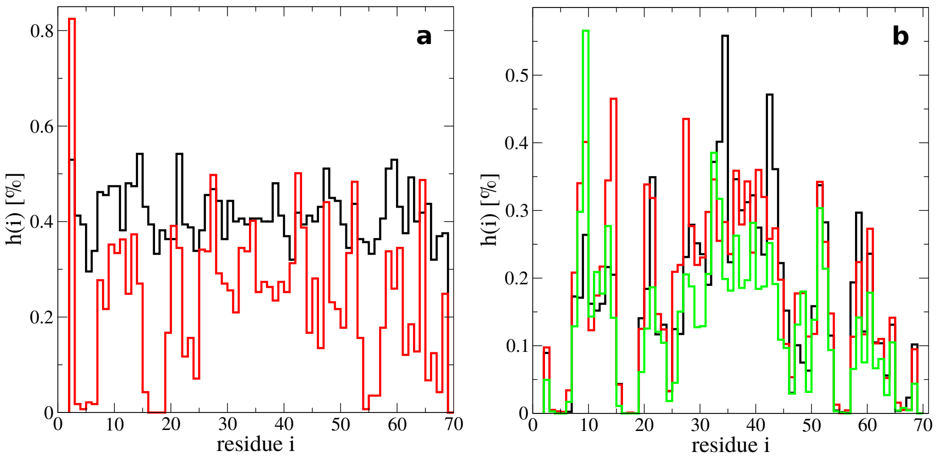


Figure S6: Percentage of helical segments per residue, $h(i)$, for ACTR determined from simulations of (a) models 1 (black) and 2 (red), and (b) models 3a (black), 3b (red) and 4 (green).

The fraction of helical segments per residue, $h(i)$, for ACTR determined from simulations of models 1, 2, 3a, 3b and 4 is nearly zero (Fig. S6), since hydrogen-bonding-like interactions are not included in these models.

#### Scaling of root mean square distances

In the main text we have presented the root mean square *normalized* distance between $C_\alpha$ atoms for two residues separated by $|j-i|$ residues along the chain, $\langle d^0_{C_\alpha}(|j-i|)\rangle$, for each of the considered models. We define $\langle d^0_{C_\alpha}(m)\rangle = \sqrt{\frac{1}{N^*_{ij}}\sum^*_{i,j}\langle d^2_{C_\alpha}(i,j)\rangle / |j-i|}$, where $\sum^*_{i,j}$ is a sum over all $ij$ pairs with $|j-i|= m$, $N^*_{ij}$ is the number of such pairs, and $d^2_{C_\alpha}(i,j) = (\mathbf{r}_j - \mathbf{r}_i)^2$. The normalization (i.e., division) by $|j-i|$ has not often been considered within the IDP literature, although it is commonplace within the polymer community. This normalization is useful for visual interpretations since $\langle d^0_{C_\alpha}(m)\rangle$ is constant for a random walk and proportional to $m^{0.1}$ for a self-avoiding random walk.[3,4] For comparisons with other IDP work, Figure S7 presents the *unnormalized* root mean square distances, $\langle d_{\mathrm{C}_\alpha}(m)\rangle = \sqrt{\frac{1}{N^*_{ij}}\sum^*_{i,j}\langle d^2_{\mathrm{C}_\alpha}(i,j)\rangle}$.

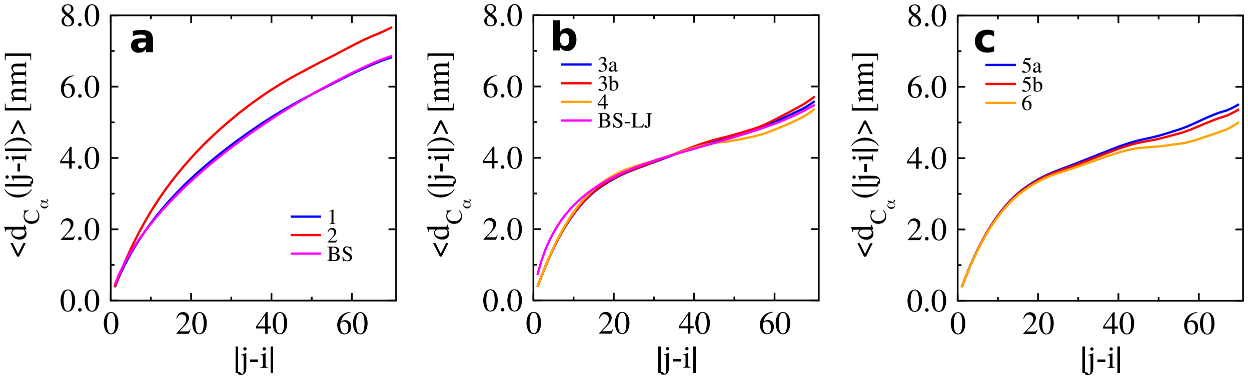


Figure S7: *Unnormalized* root mean square normalized distance between pairs of residues separated by $|j-i|$ residues along the chain, $\langle d_{\mathrm{C}_\alpha}(|j-i|)\rangle$.

In previous work,[1] these root mean square distance curves have been fit to an effective scaling law, in correspondence with the $\nu$ values presented based on the fitting of $S(q)$ in the main text. Although the results from the simulation models that behave as self-avoiding walks can be approximately fit to a single effective $\nu$ which matches that determined from $S(q)$, the other models demonstrate more complex behavior. In particular, as discussed in the main text, $\langle d^0_{\mathrm{C}_\alpha}(|j-i|)\rangle$ for the more detailed models shows three distinct regions along with three distinct effective $\nu$ values. This is an additional benefit of the normalized root mean square distance, which better highlights these regions. Figure S8 presents a plot of $\langle d^0_{\mathrm{C}_\alpha}(|j-i|)\rangle$ for various models considered in the main text, along with scaling trends to demonstrate the effective $\nu$ values at distinct separation distances.

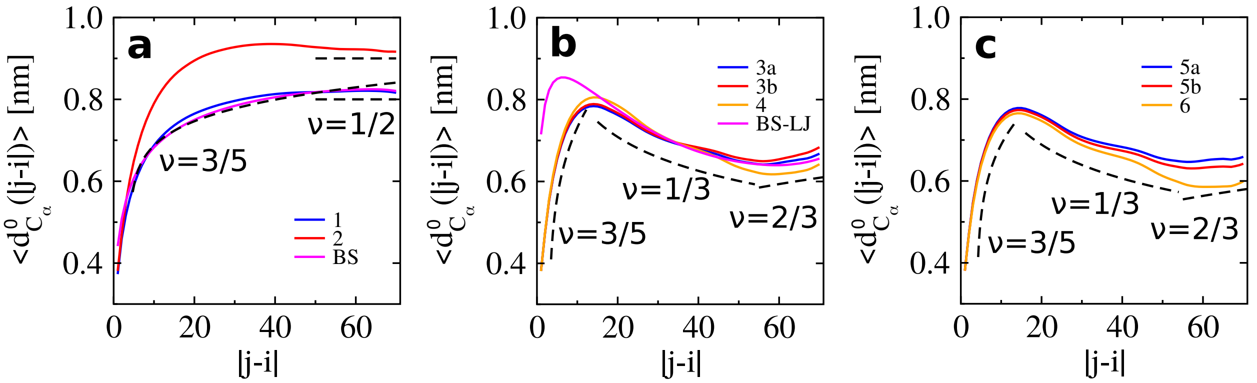


Figure S8: *Normalized* root mean square distance between pairs of residues separated by $|j-i|$ residues along the chain, $\langle d^0_{C_\alpha}(|j-i|)\rangle$. Dashed lines and labels represent effective $\nu$ values, defined with respect to the *unnormalized* root mean square distance.

#### $T$-dependence of bead-spring models

We considered two bead-spring (BS) models with WCA (BS) or Lennard-Jones (BS-LJ) potentials employed to represent interactions between monomers. The distribution of $R_g$ for these models is presented in Fig. S9. While the distribution is temperature independent for the BS model (panel (a)), $\langle R_g \rangle$ decreases with increasing temperature for the BS-LJ model.

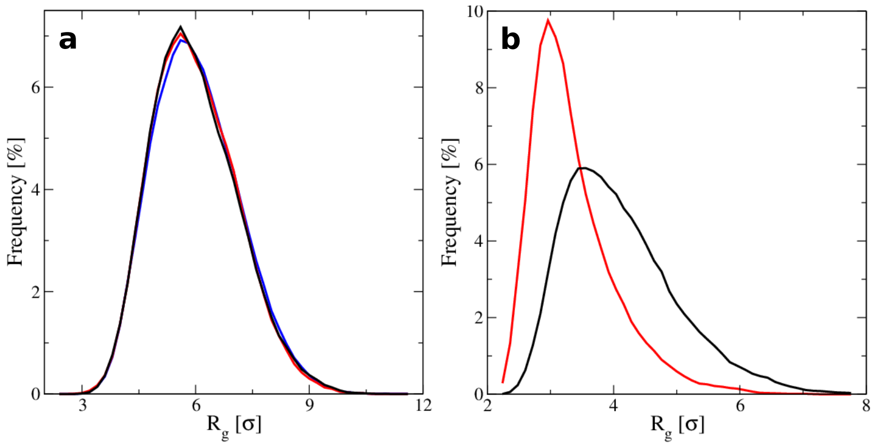


Figure S9: (a) The distribution of $R_g$ for the BS model with $N = 71$ monomers at $T = 1.5$ (blue), 2.0 (red) and 2.5 $\epsilon/k_B$ (black). (b) The distribution of $R_g$ for the BS-LJ model (with cutoff $r_c = 2.5\ \sigma$) at $T = 2.0$ (red) and 2.5 $\epsilon/k_B$ (black).

### Contact probability maps

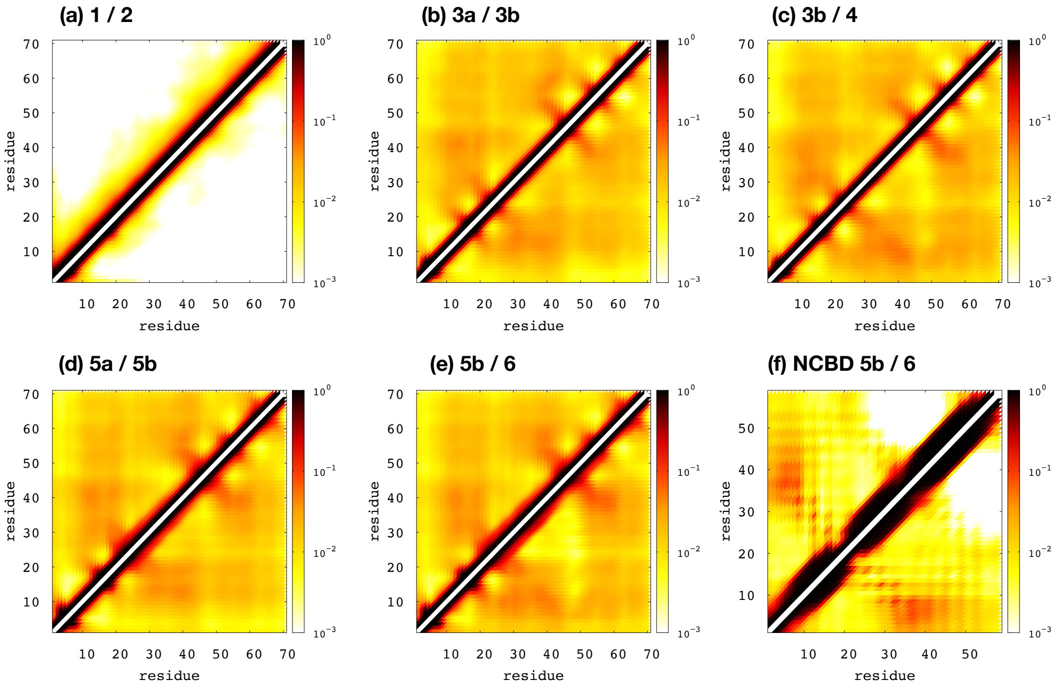


Figure S10: Heat maps of the probability that two $C_\alpha$ atoms are within a distance of 1 nm from one another. In each panel, the label "X / Y" denotes that results from model X and Y are plotted in the upper left and bottom right triangles, respectively.

### Free-energy landscapes along $R_\mathrm{g}$ and $R_\mathrm{e}$

Fig. S11 presents free-energy surfaces as a function of $R_\mathrm{g}$ and $R_\mathrm{e}$ for ACTR determined from simulations of models 3a, 3b and 4. Clearly, the shape parameters $R_\mathrm{g}$ and $R_\mathrm{e}$ do not clearly resolve the differences between the generated ensembles.

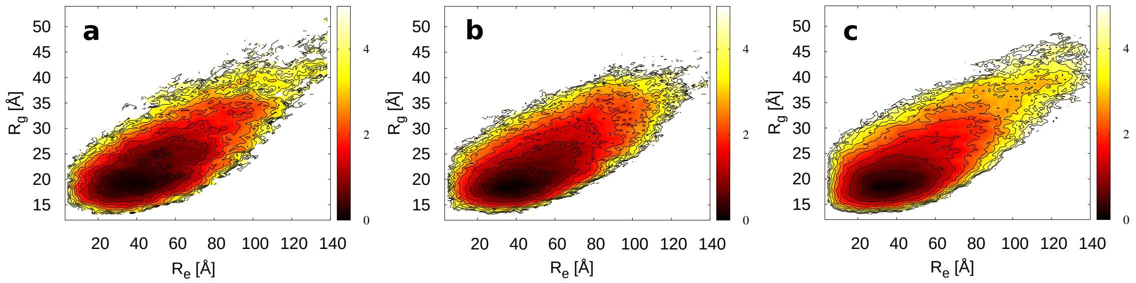


Figure S11: Free energy surfaces, in units of $k_\mathrm{B}T$, as a function of $R_\mathrm{g}$ and $R_\mathrm{e}$ for ACTR determined from simulations of models 3a (a), 3b (b) and 4 (c) at $T^*$.

#### Comparison to experimental measurements

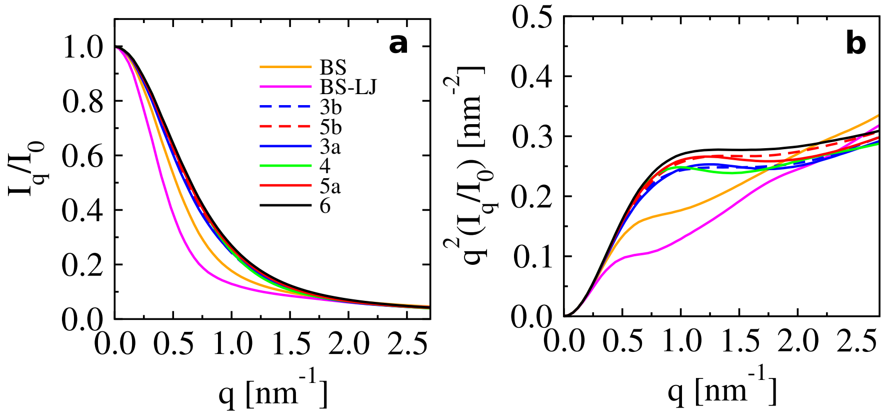


Figure S12: (a) The scattering profile $I_q/I_0$ as a function of the scattering vector $q$ for ACTR determined from simulations of the various models considered in the main text. (b) The corresponding Kratky-style plot.

Fig. S12 presents the scattering profile $I_q/I_0$ for ACTR and its corresponding Kratky representation. The results are comparable to the experimentally-determined profile, presented in Fig. 2b of.[5] The experimental profile is also presented in both forms in Figs. 1 and S1 of.[6]

#### Knots in polyA

In contrast to ACTR and polyG, we observe knotted structures in the simulations of polyA (Fig. S13). It seems that the higher rigidity (larger $h(i)$) and tight compactness (small $\langle R_{\mathrm{g}} \rangle$) of polyA allows the formation of large stable loops during the simulations, enabling threading of the termini through the loop. Knots have also been found in other polypeptides, such as in polyglutamine (polyQ).[7] In our work, the knotting probability of polyA is roughly 3%, and these knots persist for tens to thousands of $\tau$. These short lifetimes result in a small effect on the average fraction of helical segments (Fig. S14).

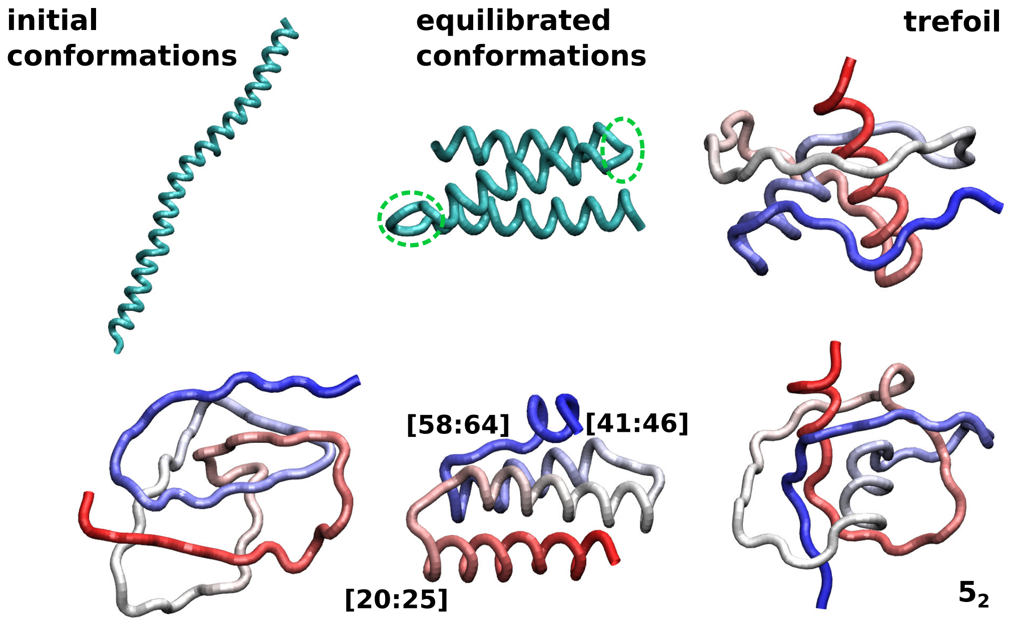


Figure S13: Representative initial conformations (left column) and equilibrated structures (middle column) for the simulations of polyA. In the middle column, the green circles denote two ranges of residues [25 : 30] and [47 : 52], where polyA is most likely to form turns, leading to smaller helicity. The bent regions of the structure in the bottom row are indicated by the numbers next to them. The right column presents representative knot structures: the trefoil and $5_2$ knot.

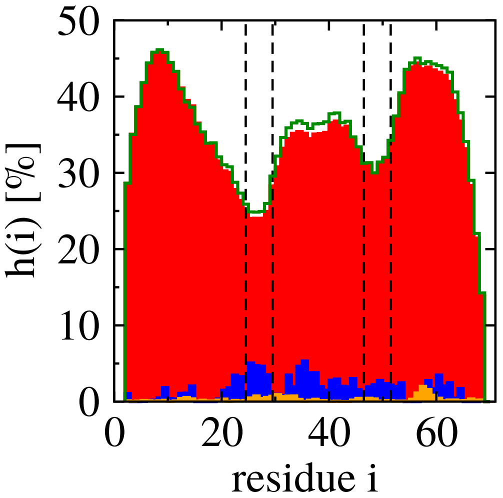


Figure S14: The percentage of helical segments per residue, $h(i)$, for polyA (red), polyG (orange) and ACTR (blue) at $T = 300$ K. The two regions marked by the black lines include the residues range from [25:30] and [47:52], and the green line indicates the results after removing the knotted frames in the polyA simulations.
